# A ciliary BBSome-ARL-6-PDE6D pathway trafficks RAB-28, a negative regulator of extracellular vesicle biogenesis

**DOI:** 10.1101/715730

**Authors:** Jyothi S. Akella, Stephen P. Carter, Fatima Rizvi, Ken C.Q. Nguyen, Sofia Tsiropoulou, Ailís L. Moran, Malan Silva, Breandán N. Kennedy, David H. Hall, Maureen M. Barr, Oliver E. Blacque

## Abstract

Cilia both receive and send information, the latter in the form of extracellular vesicles (EVs). EVs are nano-communication devices that cells shed to influence cell, tissue, and organism behavior. Mechanisms driving ciliary EV biogenesis and environment release are almost entirely unknown. Here, we show that the ciliary G-protein RAB28, associated with human autosomal recessive cone-rod dystrophy, negatively regulates EV levels in the sensory organs of *Caenorhabditis elegans*. We also find that sequential targeting of lipidated RAB28 to periciliary and ciliary membranes is highly dependent on the BBSome and PDE6D, respectively, and that BBSome loss causes excessive and ectopic EV production. Our data indicate that RAB28 and the BBSome are key *in vivo* regulators of EV production at the periciliary membrane. Our findings also suggest that EVs control sensory organ homeostasis by mediating communication between ciliated neurons and glia, and that defects in ciliary EV biogenesis may contribute to human ciliopathies.

## INTRODUCTION

Cilia are conserved microtubule (MT)-based organelles that extend from the surfaces of most eukaryotic cell types. Cilia serve a variety of functions that include motility and signal transduction, as well as the capacity to shed extracellular vesicles (EV) (Wang & Barr 2018; Carter & Blacque 2019). Most non-dividing mammalian cells possess a single, non-motile primary cilium, which acts to organize multiple signaling pathways, including those related to sonic hedgehog and PDGFα, phototransduction and olfaction (Nachury & Mick 2019). Defects in primary cilia lead to a wide range of monosymptomatic and pleiotropic human disorders (ciliopathies), frequently affecting many different tissues and organs, such as autosomal dominant polycystic kidney disease (ADPKD), various eye disorders (retinitis pigmentosa, cone-rod dystrophy) and Bardet-Biedl syndrome (BBS) (Waters & Beales 2011).

As compartmentalized and compositionally distinct organelles that lack protein synthesis machinery, cilia depend on several modes of intracellular transport to establish and control their molecular make-up (Jensen & Leroux 2017). The most well studied is intraflagellar transport (IFT), which operates bidirectionally along the ciliary MTs, driven by kinesin-II anterograde (ciliary base to tip) and cytoplasmic dynein 2 retrograde (ciliary tip to base) motors (Rosenbaum & Witman 2002). A number of cargo adapters enable IFT to transport structural and signaling proteins into and out of cilia (Lechtreck 2015; Nachury 2018). These adaptors include the IFT-A and B complexes (Mukhopadhyay et al. 2010; Liem et al. 2012; Bhogaraju et al. 2013), tubby family proteins (Badgandi et al. 2017) and the BBSome (Lechtreck et al. 2009; Jin et al. 2010). The BBSome, an octameric complex (BBS1/2/4/5/7/8/9 and BBIP10) whose assembly is regulated by the small G-protein ARL6 (Liew et al. 2014), regulates ciliary signaling cascades by removing various membrane proteins from cilia via retrograde IFT (Liu & Lechtreck 2018; Ye et al. 2018). Lipidated peripheral membrane proteins concentrate in cilia by Lipidated protein Intraflagellar Targeting (LIFT) (Jensen & Leroux 2017). In this pathway, myristoylated and prenylated proteins such as NPHP3 and INPP5E, respectively, are shuttled into cilia by prenyl-binding PDE6D (PrBPδ) and myristoyl-binding UNC119a/b, and subsequently displaced from these solubilising factors via the small G-protein ARL3 (Ismail et al. 2012; Fansa et al. 2016).

In addition to its role as a signalling hub, the cilium is also an evolutionarily conserved site of extracellular vesicle (EV) biogenesis and shedding (Wood et al. 2013; J. Wang et al. 2014; Salinas et al. 2017). EVs are highly heterogeneous by size and function, with distinct biogenesis mechanisms (Maas et al. 2017). Exosomes (typically 50-150nm) are endosomal in origin, deriving from intralumenal vesicles of a multivesicular body (MVB), which fuses with the plasma membrane to release its EV content (Raposo & Stoorvogel 2013). In contrast, ectosomes (also termed microvesicles) are larger (typically 150-500nm), and bud directly from the plasma membrane (Meldolesi 2018). EVs released from cilia appear to be ectosomal in origin (Wood et al. 2013). Cilia also harbor components components of the ESCRT pathway, which have been implicated in both ectosome and exosome biogenesis (Diener et al. 2015). Several functions have been proposed for ciliary EVs, including mediation of signalling between different cells and/or organisms (J. Wang et al. 2014; Cao et al. 2015), a waste disposal system for ciliary signalling molecules (Nager et al. 2017), a means to regulate ciliary resorption and disassembly (Long et al. 2016; Phua et al. 2017) and, in the case of the photoreceptor outer segment, material for the construction of an expanded and elaborate ciliary membrane (Salinas et al. 2017). Defects in EV biogenesis and release are associated with ciliary dysfunction indicating that EVs are an important aspect of ciliary function (Wang et al. 2015; Maguire et al. 2015; J. Wang et al. 2014; O’Hagan et al. 2017; Silva et al. 2017; Dilan et al. 2018). Indeed, EV phenotypes are a characteristic of ciliopathies such as Polycystic Kidney Disease (PKD), polycystic liver disease, and retinal degeneration (Hogan et al. 2009; Salinas et al. 2017; Dilan et al. 2018; Masyuk et al. 2010). Despite their potential importance to homeostasis and function, mechanisms driving ciliary EV biogenesis and shedding are poorly understood. Very little is known about EV cargo sorting, formation, or function, largely because their typically small size escapes detection by light microscopy. Hence, we use an *in vivo* system to study EV biogenesis in a living animal.

The nematode *C. elegans* is an excellent model for the investigation of cilium formation, specialization, transport, and function. In *C. elegans,* sensory cilia are present at the dendritic tips of sensory neurons, with most housed within sensory organs (sensilla) having environmentally exposed cuticular pores derived from surrounding glial cell processes. Of the 302 neurons that comprise the hermaphrodite nervous system, 60 are ciliated (Ward et al. 1975; Perkins et al. 1986; Doroquez et al. 2014). *C. elegans* males possess an additional 54 ciliated neurons not present in the hermaphrodite (Sulston et al. 1980; Cook et al. 2019; Molina-García et al. n.d.). *C. elegans* cilia possess a similar basic structural plan, extending from a degenerate basal body within a bulge in the dendritic ending termed the periciliary membrane compartment (PCMC), which acts as a trafficking hub for sorting proteins between ciliary and cell body destinations (Kaplan et al. 2012; Nechipurenko et al. 2017; Serwas et al. 2017). Axonemes that extend from the PCMC of the majority of *C. elegans* cilia can be divided into transition zone (TZ), middle and distal segments which display tremendous diversity in terms of their MT arrangements between different cell-types (Perkins et al. 1986; Doroquez et al. 2014). The most well-studied cilia types in *C.elegans* are the amphid and phasmid cilia that perform a wide array of sensory functions including chemosensation, osmosensation and thermosensation (Bargmann 2006).

In the brain of the worm, the cephalic male (CEM) and shared inner labial (IL2) neurons are the only known ciliated EV-releasing neurons (EVNs) as defined by imaging of fluorescently-tagged EV cargos released outside the animal (J. Wang et al. 2014). These EV-releasing cilia display non-canonical variations of the 9+0 doublet microtubule axoneme. During sexual maturation, CEM doublets splay to 18 A-tubule and B-tubule singlets while the IL2 transition zone remodels from 9+0 to 4-7+0 (Perkins et al. 1986; Doroquez et al. 2014; Silva et al. 2017; Akella et al. 2019). The tubulin code via tubulin glutamylation and tubulin isotype control this unique CEM ciliary ultrastructure (Silva et al. 2017; O’Hagan et al. 2017). Environmentally-released EVs contain cargoes, including the polycystin PKD-2, that regulate mating behaviors and, thus, their bioactivity promotes inter-animal communication (J. Wang et al. 2014; Wang et al. 2015; Maguire et al. 2015; Silva et al. 2017). In *C. elegans*, EVs are ‘shed’ at the PCMC and retained within pores of the cephalic and inner labial sensory organs as visualized by transmission electron microscopy (TEM) and tomography reconstructions; EVs are also ‘released’ to the animal’s external environment as observed by fluorescently-tagged EV cargos such as PKD-2::GFP (J. Wang et al. 2014). Ciliary EV release is controlled by the tubulin code writers (a glutamylase TTLL-11 and a- tubulin TBA-6), erasers (deglutamylase CCPP-1) and readers (the IFT machinery and kinesin-3 KLP-6) (J. Wang et al. 2014; Wang et al. 2015; Maguire et al. 2015; Silva et al. 2017; O’Hagan et al. 2017). In mutants of these genes, EVs are shed and accumulate in pores of sensory organs but are not released into the environment. How the cilium controls the site of EV shedding (ciliary base versus ciliary tip) or EV environmental release is not known.

A number of small G-proteins, including those of the Rab family, regulate ciliogenesis and cilia functions, as well as EV biogenesis (Li & Hu 2011; Blacque et al. 2018; Knödler et al. 2010; Blanc & Vidal 2018; Meldolesi 2018). Here we focus on Rab28, which is associated with autosomal recessive cone-rod dystrophy (arCRD) (Roosing et al. 2013; Riveiro-Álvarez et al. 2015; Lee et al. 2017). A similar photoreceptor degeneration is found in many ciliopathies, including Bardet-Biedl syndrome (Waters & Beales 2011). In *C. elegans* amphid and phasmid cilia, RAB-28 is a BBS-8-dependent IFT cargo that associates with the periciliary membrane (PCM) when in a GTP-bound conformation (Jensen et al. 2016). Functionally, *C. elegans* RAB-28 disruption leads to defects in amphid pore formation via a cell non-autonomous mechanism previously speculated to involve EVs (Jensen et al. 2016). An EV-related function for RAB-28 is also suggested by its significant overrepresentation in *C. elegans* EVN transcriptomes (Wang et al. 2015).

Here we dissect the transport pathway that regulates RAB-28 localization in amphid and EVN cilia, and find that its PCM association is dependent on the BBSome and, to a lesser extent, its regulator ARL-6, whilst its IFT transport requires LIFT components PDL-1 (PDE6D) and ARL-3. We also identify cell-specific distinctions in how RAB-28 transport machinery is deployed, with PDL-1 dispensable for maintaining RAB-28’s PCM levels in amphid/phasmid neurons but not EVNs. Functional analyses revealed that BBSome mutants phenocopy the enlarged amphid pores observed in RAB-28-disrupted worms. Furthermore, loss of RAB-28 or BBSome genes, but not PDL-1, causes ectopic EV accumulation in the cephalic sensory organ pore, suggesting that RAB-28 and the BBSome negatively regulate ciliary EV biogenesis and/or shedding at the PCM. We conclude that RAB-28 and the BBSome function at the PCM to regulate ciliary EV production, with defects resulting in excessive EV shedding and accumulation within sensory organs. This research reveals a BBSome-ARL-6-PDL-1 network for targeting RAB-28, a critical EV regulator, to specific ciliary membranes, thus restricting the production of EVs to distinct sites. The members of this network as well as the EV cargo PKD-2 are well established ciliary proteins associated with several human genetic diseases (Waters & Beales 2011; Reiter & Leroux 2017) and, as such, dysregulated ciliary EV production may be a factor in the pathology of certain ciliopathies.

## METHODS

### *C. elegans* strains and maintenance

All strains were cultured according to standard protocols (Brenner 1974). Briefly worms were cultured on plates of nematode growth medium (NGM), seeded with a lawn of OP50 *E. coli.* Plates were incubated at either 20°C or 15°C to slow development. Reporter strains were crossed into mutant backgrounds and double mutants generated by standard crossing strategies. Mutations were followed by genotyping PCR.

#### STRAIN LIST

OEB803 N2; *oqEx304*[*gfp::rab-28^Q95L^ + unc-122p::gfp*]

OEB972 *arl-6(ok3472); oqEx304*[*gfp::rab-28^Q95L^ + unc-122p::gfp*]

OEB970 *bbs-5(gk507); oqEx304*[*gfp::rab-28^Q95L^ + unc-122p::gfp*]

OEB805 *bbs-8(nx77); oqEx304*[*gfp::rab-28^Q95L^ + unc-122p::gfp*]

OEB806 *pdl-1(gk157); oqEx304*[*gfp::rab-28^Q95L^ + unc-122p::gfp*]

OEB807 *arl-3(tm1703)*; *oqEx304*[*gfp::rab-28^Q95L^ + unc-122p::gfp*]

OEB971 *pdl-1(gk157); bbs-8(nx77); oqEx304*[*gfp::rab-28^Q95L^ + unc-122p::gfp*]

OEB973 *him-5(e1490); oqEx304*[*gfp::rab-28^Q95L^ + unc-122p::gfp*]

OEB974 *arl-6(ok3472); him-5(e1490); oqEx304*[*gfp::rab-28^Q95L^ + unc-122p::gfp*]

OEB975 *bbs-5(gk507); him-5(e1490); oqEx304*[*gfp::rab-28^Q95L^ + unc-122p::gfp*]

OEB976 *bbs-8(nx77) him-5(e1490); oqEx304*[*gfp::rab-28^Q95L^ + unc-122p::gfp*]

OEB977 *pdl-1(gk157); him-5(e1490); oqEx304*[*gfp::rab-28^Q95L^ + unc-122p::gfp*]

OEB978 *pdl-1(gk157); arl-6(ok3472); him-5(e1490); oqEx304*[*gfp::rab-28^Q95L^ + unc-122p::gfp*]

PT621 *him-5(e1490); myIs4* [*PKD-2::GFP+Punc-122::GFP*]

PT2679 *him-5(e1490); myIs23*[*cil-7p::gCIL-7::GFP_3’UTR+ccRFP*]

PT3189 *rab-28(tm2636); him-5* OEB945 *arl-6(ok3472); him-5* OEB947 *bbs-8(nx77); him-5*

PT2984 *rab-28(tm2636);him-5; myIs4* [*PKD-2::GFP+Punc-122::GFP*]

PT3265 *rab-28(tm2636);him-5; myIs23*[*cil-7p::gCIL-7::GFP_3’UTR+ccRFP*]

PT3190 *pha-1(e2123); him-5; myEx905*[*rab-28p::sfGFP+PBX*]

PT3356 *pha-1(e2123); him-5; myIs20* [*klp-6p::tdtomato*] *myEx905*[*rab-28p::sfGFP+PBX*]

OEB948 *rab-28(gk1040); him-5 myIs23*

OEB949 *arl-6(ok3472);him-5 myIs23*

OEB951 *pdl-1(gk157); him-5 myIs23*

### Fluorescence microscopy

For PKD-2::GFP, CIL-7::GFP, *rab-28p*::sfGFP and *klp-6p:*:tdTomato (in Figures 3 - 5), L4 males were isolated the previous day to provide virgin adults for imaging the next day. Males were placed on 4% agarose pads, and immobilized in 10mM levamisole before imaging. For all CIL-7::GFP imaging (except that for comparison of wild-type and *bbs-8* mutants), males were let to settle down in levamisole for 7-8 minutes prior to imaging. Epifluorescence imaging was performed using an upright Zeiss Axio Imager D1m. Images were acquired using a digital sCMOS camera (C11440-42U30, Hamamatsu Photonics). The microscope was controlled by Metamorph 7.1 to acquire Z stacks. All images were analyzed using FIJI.

To compare average fluorescence intensities of CIL-7::GFP and PKD-2::GFP between wild-type and mutant males, lines were traced across the CEM cilium and the plot profile function on FIJI was used to generate intensity values across several points of the cilium. These numbers were then averaged to give an intensity value for each cilium across many animals. To obtain line graphs depicting differences in the fluorescence intensity values across different points along the CEM cilia, cilia were traced from the tip to PCMC and the intensity levels at different points along the cilium were obtained for several animals. The intensity of each point along the cilium was then averaged across several cilia/ animals. Only the points that had intensity values across all samples examined were plotted in the line graph.

For EV particle quantification, EVs containing PKD-2::GFP were counted from blinded images of age matched wild-type and mutant adult males expressing *myIs4*[PKD-2::GFP] as described previously (J. Wang et al. 2014). EVs containing CIL-7 were counted from images of age matched wild-type and mutants expressing *myIs23*[CIL-7::GFP]. EV particles were quantified from Z projections using the ROI manager tool on FIJI.

Statistical analysis: Raw data was sorted and arranged using Microsoft Excel. Statistical analyses were done using GraphPad Prism V5. We used standard symbols to depict whether the P values were statistically significant (* for p<0.05, ** for p<0.005, and *** for p< 0.0005).

For analysis of RAB-28^Q95L^ localization, hermaphrodite or male worms were placed on 5% agarose pads on glass slides and immobilized with 40 mM levamisole. L4 males were isolated the previous day to provide virgin males for imaging. Epifluorescence imaging was performed on an upright Zeiss AxioImager M1 microscope with a Retiga R6 CCD detector (Teledyne QImaging). Confocal imaging was performed on an inverted Nikon Eclipse Ti microscope with a Yokogawa spinning-disc unit (Andor Revolution) and images were acquired using an iXon Life 888 EMCCD detector (Andor Technology). All image analysis was performed using Fiji (Schindelin et al. 2012). For kymography, time-lapse (multi tiff) movies of IFT along cilia were taken at 250 ms exposure and 4 fps. Kymographs were generated from multi tiff files using the KymographClear ImageJ plugin (Mangeol et al. 2016).

### Electron microscopy

High pressure freeze fixation (HPF) and freeze substitution for TEM on CEM cilia: wild-type CB1490 (*him-5*), PT3189 (*rab-28(tm2636); him-5*), OEB947 (*bbs-8(nx77); him-5*) were collected as L4 males the day before freeze fixation to provide virgin day 1 adult males on the day of fixation. Animals were subjected to high-pressure freeze fixation using a HPM10 high-pressure freezing machine (Bal-Tec, Switzerland). Animals were slowly freeze substituted in 2% osmium tetroxide, 0.2% uranyl acetate and 2% water in acetone using RMC freeze substitution device (Boeckeler Instruments, Tucson, AZ, USA) (Weimer 2006). Samples were infiltrated with Embed 812 resin over 3 days prior to embedding in blocks.

Most animals were collected in 70 nm-thick plastic serial sections collected on copper slot grids and were post-stained with 2% uranyl acetate in 70% methanol, followed by washing and incubating with aqueous lead citrate. TEM images were acquired on either a Philips CM10 transmission electron microscope operating at 80 kV or a JEOL JEM-1400 transmission electron microscope operating at 120 kV.

TEM of amphid pores of BBS mutants (in Figure 2) was performed as previously described (Sanders et al. 2015). Briefly, hermaphrodites were fixed overnight at 4°C in 2.5% glutaraldehyde in Sørensen’s phosphate buffer (0.1M, pH7.4). Samples were post-fixed in 1% osmium tetroxide and dehydrated in an ascending gradient series of ethanol concentrations prior to Epon 812 resin embedding overnight. Subsequently, 90 nm sections were cut using a Leica EM UC6 Ultramicrotome, collected on copper grids and post-stained with 2% uranyl acetate and 3% lead citrate. Imaging was performed on a Tecnai T12 (FEI) using an accelerated voltage of 120 kV. Electron tomography was performed as described in (Silva et al. 2017). Slice views of tomograms on CEM ciliary tips were exported as tiff files.

For quantification of EVs using TEM: All quantifications of EV numbers were done from serial section TEM images of males of WT and *rab-28(tm2636)* animals. For comparison of EV numbers in the sub-distal region of the cephalic sensory organ lumen, EVs were counted between the regions where the CEM cilia have 18 singlet MTs up to the anterior-most section where CEM and CEP cilia share the lumen. For comparison on EV numbers at the TZ level of CEM cilia in cephalic sensory organs: EVs were counted in the cephalic lumen between the region where all the TZ microtubules were doublets up to the region where all the TZ microtubules terminated. For comparison on EV numbers at the PCMC level: EV numbers were counted 140 nm posterior to the region after the TZ microtubules terminated. To account for whether the EV phenotypes in *rab-28* were due to a specific defect, and not to changes in the positioning of CEM cilia within the cephalic lumen, the CEP and OLQ cilia were also used as landmarks while scoring EV numbers.

## RESULTS

### RAB-28 transport to and within amphid and phasmid cilia is regulated by the BBSome, ARL-6 and LIFT-associated PDL-1

In amphid and phasmid neurons, we previously showed that the IFT motility and periciliary membrane (PCM) targeting of GFP-tagged RAB-28^Q95L^, a putative GTP-preferring and active form of the G-protein, is dependent on the BBSome component *bbs-8* (Jensen et al. 2016). To further establish the role of the BBSome in RAB-28 ciliary localization and transport we assessed GFP-RAB-28^Q95L^ in *bbs-5* null mutant cilia. We also assessed GFP-RAB-28^Q95L^ localization in a null mutant of *arl-6*, which regulates membrane-targeting of the BBSome in mammalian cells (Jin et al. 2010). We found that GFP-RAB-28^Q95L^ mislocalization in the amphid and phasmid cilia of *bbs-5* worms is indistinguishable from *bbs-8* mutants; GFP signals are diffuse throughout the neurons, with no detectable PCM enrichment or IFT movement (Figure 1; enhanced contrast images and expressivity shown in Supplementary Figure 1A and C). By contrast, *arl-6* mutants display a weaker phenotype, with modestly reduced levels of RAB-28 at the PCM and a diffuse dendritic localization (Figure 1). Also, in *arl-6* phasmid cilia, the frequency of GFP-RAB-28^Q95L^-positive IFT trains is increased twofold (Supplementary Figure 1B and Supplementary Movie 1). These data indicate that a complete and properly regulated BBSome is required for normal targeting and retention of RAB-28 to the PCM and that ARL-6 inhibits RAB-28 association with IFT trains, at least in amphid and phasmid neurons.

**Figure 1.**
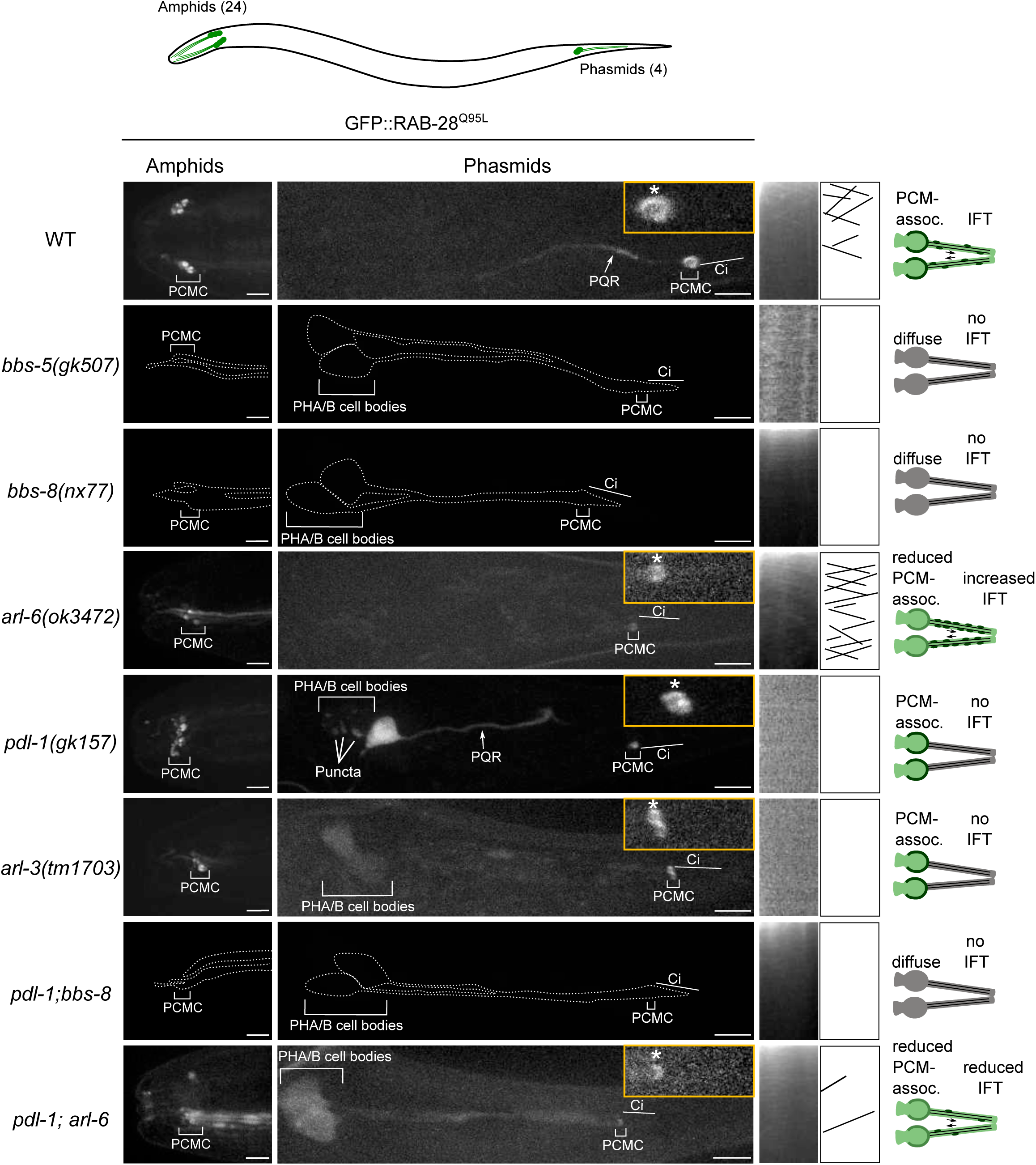
A BBSome-ARL-6-PDL-1 network targets RAB-28 to sensory cilia. Representative confocal z-projection images of amphid (head) and phasmid (tail) neurons from hermaphrodites expressing GFP::RAB-28^Q95L^. Anterior is to the left and all images taken at identical exposure settings. Traced outlines in *bbs-5*, *bbs-8* and *pdl-1;bbs-8* panels are derived from intensity-adjusted images (see Supplementary Figure 1). Insets; higher magnification images of phasmid cilia, with PCMC denoted by asterisks. Kymograph x-axis represents distance and y-axis time (scale bars; 5 s and 1 μm), and both retrograde and anterograde particle lines are shown in the kymograph schematics. Schematics on the right summarize the phenotypes observed in a pair of phasmid cilia. Scale bars; 10 μm. PCMC: periciliary membrane compartment; ci: cilium; PQR: additional ciliated neuron in the tail that occasionally expresses the RAB-28 reporter.

As RAB-28 is a prenylated protein, we investigated whether RAB-28 requires lipidated intraflagellar transport (LIFT) for its ciliary localization, using null mutants of *arl-3* and *pdl-1*, the *C. elegans* orthologs of *ARL3* and *PDE6D*. We found that whilst the PCM localization of GFP-RAB-28^Q95L^ remains intact, the reporter’s IFT motility is completely lost in these mutants (Figure 1 and Supplementary Figure 1C). An additional punctate localization was also detected for GFP-RAB-28^Q95L^ in the phasmid cell bodies of *pdl-1* mutants. Thus, like the BBSome, LIFT is also required for RAB-28 association with IFT trains in amphid and phasmid neurons, although it is dispensable for RAB-28 targeting to the PCM of these cells.

To explore the genetic relationship between the BBSome, ARL-6 and LIFT-dependent targeting mechanisms, we assessed RAB-28 localization in *pdl-1;bbs-8* and *pdl-1;arl-6* double mutants. GFP::RAB-28^Q95L^ localization and IFT behavior in *pdl-1;bbs-8* mutants is identical to that of *bbs-8* single mutants, reproducing the complete loss of RAB-28’s PCM association and IFT in that strain (Figure 1 and enhanced contrast images in Supplementary Figure 1A), indicating that *bbs-8* is epistatic to *pdl-1*. Similarly, in *pdl-1;arl-6* worms, the reduced GFP::RAB-28^Q95L^ PCM localization phenocopies that of the *arl-6* single mutant, although there are some additional diffuse signals in the cilium and distal dendrite (Figure 1). With regard to RAB-28’s IFT behavior phenotype, *pdl-1* loss strongly suppresses the increased IFT frequency phenotype associated with *arl-6* disruption, further supporting the critical role of PDL-1 in regulating RAB-28 loading onto IFT trains (Supplementary Figure 1B and C and Supplementary Movie 1). It is notable, however, that a low level of RAB-28 IFT movement remains in at least some *pdl-1;arl-6* cilia (Supplementary Figure 1B and C and Supplementary Movie 1). Since detectable IFT was never observed in the *pdl-1* single mutant, the latter data could indicate that *arl-6* and *pdl-1* mutations are mutually suppressive with respect to RAB-28’s IFT motility phenotype, consistent with an opposing functional relationship between these genes in regulating the formation of RAB-28-positive IFT trains.

Taken together, these data reveal a ciliary trafficking pathway in amphid and phasmid neurons whereby activated RAB-28 is initially targeted to the PCM by the BBSome, and to a lesser extent ARL-6, and subsequently solubilized by PDL-1 for loading onto IFT trains, a step inhibited by ARL-6.

### *C. elegans* BBS mutants phenocopy the amphid pore defects of RAB-28-disrupted worms

The amphid sensory organ or sensillum contains 10 rod-like cilia (from 8 neurons) located in an environmentally-exposed channel, or ‘pore’, formed by the dendrites and ciliary axonemes punching through surrounding amphid sheath and socket glial cell processes (Ward et al. 1975; Doroquez et al. 2014). Previously we showed that overexpression of GTP or GDP-preferring RAB-28 cell non-autonomously induces defects in amphid pore formation and the surrounding sheath cell (Jensen et al. 2016). Given the requirement of the BBSome and ARL-6 for ciliary localization of RAB-28, we investigated amphid pore structure in chemically-fixed BBS gene mutants using transmission electron microscopy (TEM). We found that *bbs-5* and *bbs-8* mutant hermaphrodites display similar defects to hermaphrodites overexpressing RAB-28^T26N^ (GDP-preferring), namely large deposits of electron dense material in the sheath cell process at all levels surrounding the cilia (Figure 2A). Additionally, much like worms overexpressing RAB-28^Q95L^ (GTP-preferring) (Jensen et al. 2016), *bbs-5* and *bbs-8* mutant hermaphrodites exhibit highly distended amphid pores, although with much darker staining of the pore’s extracellular matrix (ECM) (Figure 2A and B). Also, at the level of the ciliary middle segments and transition zones, where the pore is enlarged, *bbs-5* and *bbs-8* mutant pores contain 11 cilia, instead of the usual 10 (Figure 2B and C). A weaker phenotype was observed in the *arl-6* mutant; whilst there are electron dense deposits in the sheath cell at the middle segment, transition zone and PCMC levels of the pore, the size and ECM density of the pore is normal (Figure 2A). Together, these data show that disruption of the BBSome, or ARL-6 to a lesser extent, phenocopies the amphid pore and sheath cell defects observed in *rab-28*-disrupted hermaphrodites. As with *rab-28*, *C. elegans* BBS genes are expressed exclusively in ciliated neurons (Ansley et al. 2003; Fan et al. 2004), indicating that their effects on the amphid sheath cell and pore are also due to cell non-autonomous functions. Thus, in the amphid sensory organ, the BBSome, ARL-6 and RAB-28 function together in common cell non-autonomous pathways.

**Figure 2.**
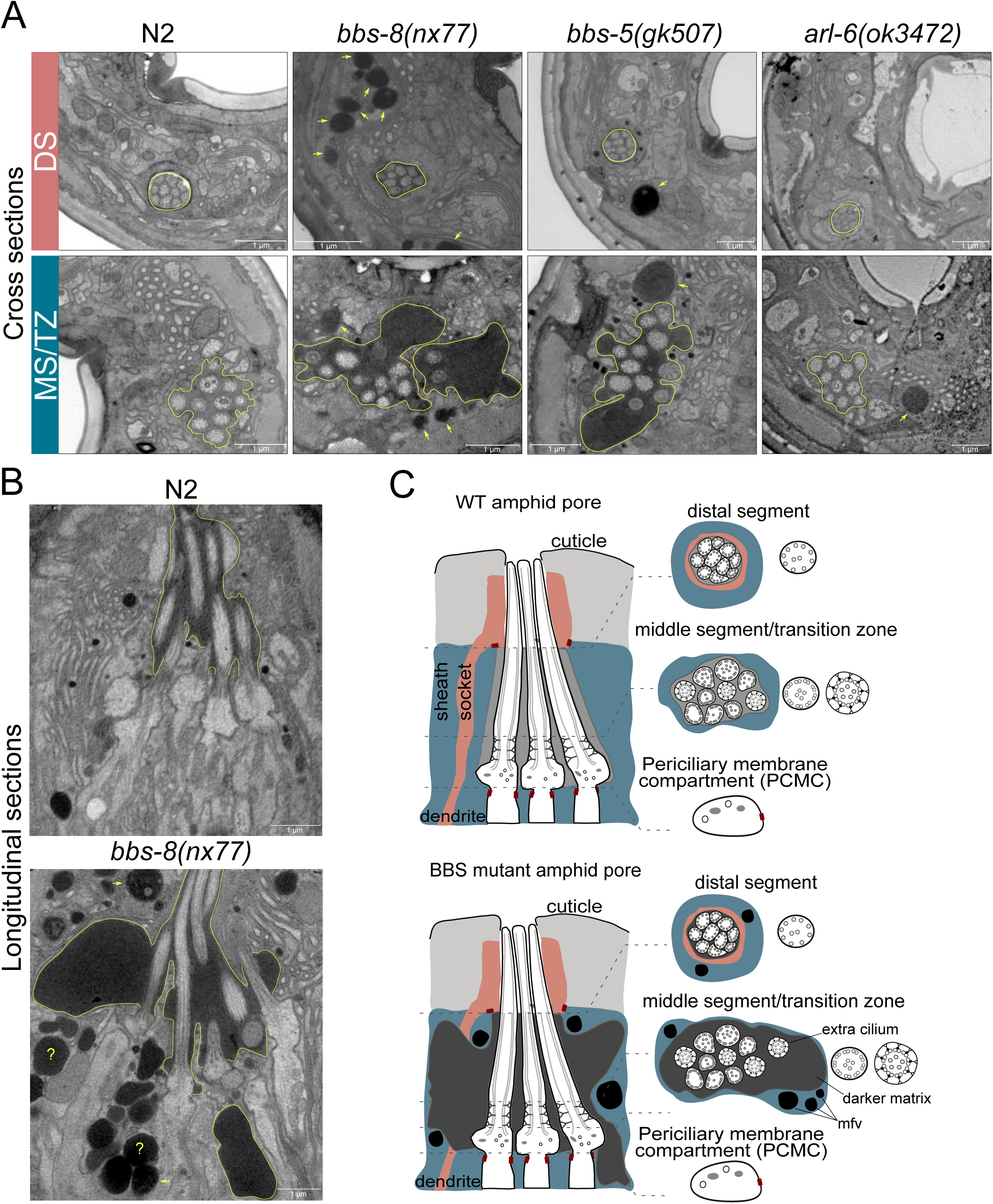
BBSome and *arl-6* mutant hermaphrodites display defects in amphid sensory pore structure and/or associated glia. **(A)** Transmission electron microscopy (TEM) images showing the amphid channels of N2 (WT), *bbs-5*, *bbs-8* and *arl-6* mutants in cross section, at the positions of ciliary distal segments (DS) and middle segments (MS)/transition zones (TZ). Arrows point to matrix-filled vesicles (mfv) within the cytoplasm of the sheath glial cell. The extracellular matrix-filled amphid pore volume is also highlighted. Scale bars; 1μm (all panels). **(B)** TEM images of longitudinal sections through the amphid cilia of N2 (WT) and *bbs-8* mutant worms, highlighting the expanded pore volume and accumulated mfv in the sheath cell (arrows). Question marks denote densities for which identification as either an mfv or an expanded pore region is ambiguous. Scale bars; 1μm. **(C)** Cartoon representations of amphid pore ultrastructure in WT and BBSome mutant worms, in longitudinal and cross section. Only three cilia are shown in the longitudinal schematic for simplicity.

### A modified BBSome-ARL-6-PDL-1 network targets RAB-28 to the periciliary membrane and cilia of extracellular vesicle-releasing neurons (EVNs)

Previously we speculated that the cell non-autonomous roles of *rab-28* could be related to a ciliary extracellular vesicle (EV) pathway based on our finding that *rab-28* is one of 335 genes whose expression is enriched in EV releasing neurons (EVNs) (Wang et al. 2015; Jensen et al. 2016). Therefore, we sought to examine possible EV-related roles for RAB-28 in the cephalic male CEM cilia for which we have identified an established repertoire of cargoes, EV bioactivity assays, known EV biogenesis and release regulators, and assayable phenotypes to test for a role in EV biology (J. Wang et al. 2014; Wang et al. 2015; Maguire et al. 2015; Silva et al. 2017; O’Hagan et al. 2017). First, however, we wished to confirm that *rab-28* was expressed in EVNs and if the encoded protein also trafficks to cilia in these neurons via a BBSome-LIFT pathway, as is the case in amphid and phasmid cells.

To confirm that *rab-28* is expressed in EVNs, we examined transgenic animals co-expressing the EVN reporter *klp-6p*::tdTomato and a transcriptional GFP reporter under the control of *rab-28* promoter (*rab-28p*::sfGFP). We found that the *rab-28* reporter is expressed in the IL2 neurons present in both males and hermaphrodites, as well as all 21 male-specific EVNs, namely the CEMs in the head and the ray RnB and hook HOB neurons in the tail (Figure 3A).

Next we explored RAB-28 subcellular localization in male-specific EVNs using our GFP::RAB-28^Q95L^ reporter. Like amphid and phasmid channel neurons, RAB-28 is enriched at the PCM of CEM and RnB neurons (Figure 3B). However, unlike amphid and phasmid cells, a pool of RAB-28 also occurs within the distal ends of the CEM and ray RnB cilia. Moving RAB-28-positive IFT particles were only relatively rarely detected in CEM and RnB cilia (Figure 3C and Supplementary Movie 2), consistent with our previous report showing that IFT in CEM cilia is less frequent than in amphid cilia (Morsci & Barr 2011). Interestingly, almost all detectable RAB-28-positive IFT trains in male EVN cilia move in the retrograde direction (Figure 3C). We did not observe RAB-28 in environmentally released EVs, indicating that RAB-28 is not EV cargo.

To address the requirement of RAB-28 transport machinery in EVNs, we generated *him-5* male mutants of the BBSome (*bbs-5*, *bbs-8)*, *arl-6* and *pdl-1* expressing the GFP::RAB-28^Q95L^ reporter. In EVN cilia, GFP::RAB-28^Q95L^ PCM localization was absent in *bbs-5* and *bbs-8* worms, whilst reduced in *arl-6* and *pdl-1* worms (Figure 3B), similar to phenotypes observed in the amphid channel and phasmid cilia in these mutants (Figure 3B). Also, the distal pool of RAB-28 in CEM and RnB cilia was reduced or absent in all of these mutants (Figure 3B). Like in amphid and phasmid neurons, IFT movement of RAB-28^Q95L^ was absent in *bbs-5, bbs-8* and *pdl-1* mutant EVNs. However, for *arl-6* worms, RAB-28^Q95L^ IFT movement was also not detectable in EVNs, which is in striking contrast to the increased frequency of RAB-28-positive IFT trains in amphid and phasmid neurons (Jensen et al. 2016). To assess the genetic relationship of *pdl-1* and *arl-6* in EVNs, we analyzed a *pdl-1;arl-6* double mutant and found that GFP::RAB-28^Q95L^ shows no PCM enrichment, with a diffuse cytoplasmic localisation throughout the neurons; thus, in contrast to amphid and phasmid neurons, *pdl-1* and *arl-6* demonstrate a synthetic relationship in targeting RAB-28 to EVN cilia.

**Figure 3.**
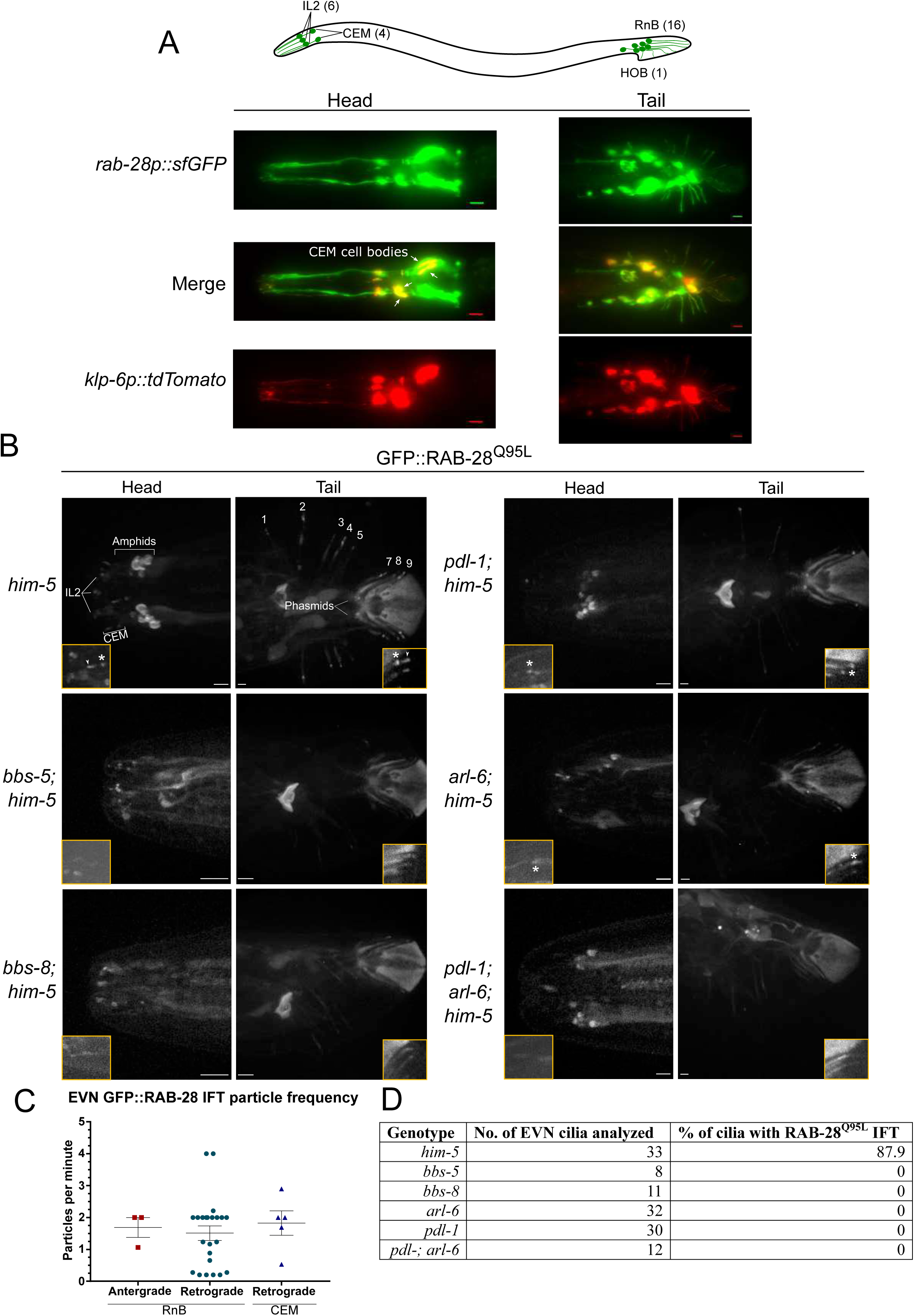
RAB-28 is expressed in and trafficked to EVN cilia via a modified BBSome-ARL-6-PDL-1 pathway. **(A)** Epifluorescence z-projections of the heads and tails of *C. elegans* males expressing sfGFP under the *rab-28* promoter and *klp-6p::tdTomato* (EVN cilia-specific reporter). Anterior is to the left. Scale bars; 5 μm. **(B)** Representative images of *him-5* (control), *bbs-5; him-5*, *bbs-8 him-5*, *pdl-1; him-5*, *arl-6; him-5* and *pdl-1; arl-6; him-5 C. elegans* males expressing GFP::RAB-28^Q95L^ in EVNs (reporter also expressed in amphid and phasmid neurons). Insets show higher magnification images of CEM (head) and RnB (numbered 1-9 in the tail) cilia. Asterisks indicate PCMC; white arrowheads indicate accumulated GFP::RAB-28^Q95L^ in the distal region of CEM and RnB cilia. Anterior is to the left. Scale bars 5 μm. **(C)** Scatter plots of GFP::RAB-28 IFT particle frequency in CEM and RnB male cilia. Error bars show SEM. Data are from 26 worms. **(D)** Table summarizing the percentage of CEM and RnB cilia with detectable IFT movement of GFP::RAB-28^Q95L^ in the indicated mutants.

Together, these data reveal that the BBSome/ARL-6/PDL-1 network that regulates RAB-28 localization and IFT behavior in amphid and phasmid neurons is also functioning in EVNs. Although similar, there are differences in how the network is deployed in these different cell types. In amphid and phasmid neurons, GTP-bound RAB-28 is enriched at the PCM and undergoes continuous IFT, which is promoted by PDL-1 and inhibited by ARL-6. In EVNs, the PCM is also a site of RAB-28^Q95L^ enrichment along with an additional RAB-28^Q95L^ pool in the distal cilium. However, RAB-28 IFT events are infrequent in this cell type and positively regulated by ARL-6.

### RAB-28 regulates the localization of EV-associated cargoes

In the male head, four quadrant cephalic sensilla contain cilia located on the distalmost dendritic endings of CEM and CEP neurons that are surrounded by socket and sheath glial support cells (Figure 4A, Figure 5A). The CEM cilium curves out and the tip protrudes to the environment via a cuticular pore, while the CEP cilium curves in and embeds in the cuticle. The cephalic lumen formed by the sheath and socket cells contains CEM-derived EVs as visualized by TEM and electron tomography (Wang et al. 2014). To determine whether *rab-28* has an EV-related role in CEMs, we started by examining the localization of fluorescently tagged ciliary and EV cargo proteins in *rab-28* mutants. We used two deletion alleles of *rab-28*, the previously studied *gk1040* null allele (998 bp deletion) (Jensen et al. 2016) and the *tm2636* allele, which is a 147 bp deletion that leads to a premature stop codon and presumable loss of *rab-28* gene function (Figure 4A).

**Figure 4.**
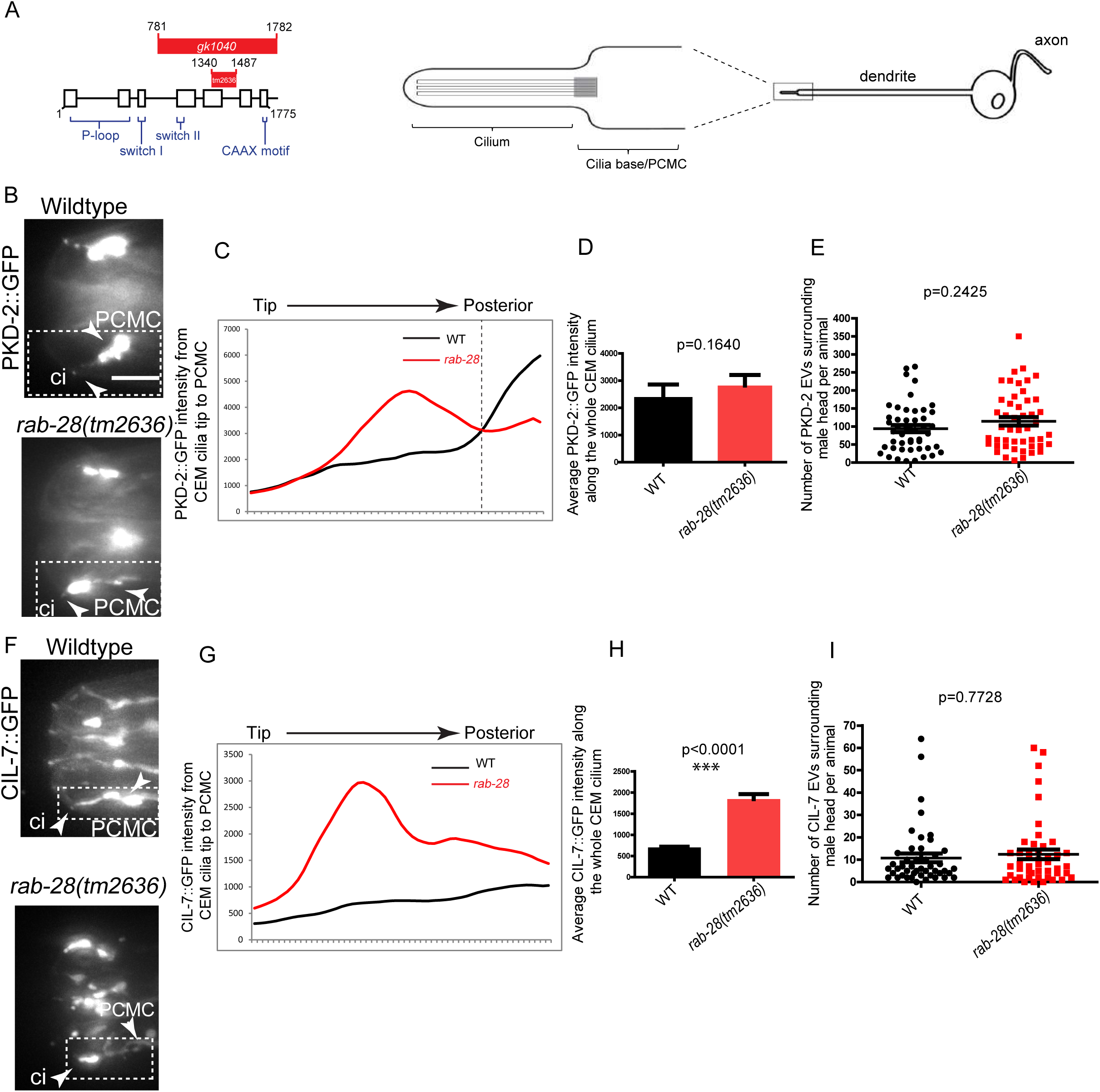
RAB-28 regulates the localization of ciliary EV cargoes in EV releasing CEM cilia. **(A)** Left: genomic structure of *rab-28* showing the location of the deleted region in *gk1040* and *tm2636* mutant alleles. Right: Cartoon reproduced from (Bae et al. 2006) depicting the morphology of the CEM neurons of *C. elegans.* CEM cilia are present at the tips of sensory dendrites. Inset shows the location of the ciliary base/ PCMC and the MT-based cilium. **(B)** Fluorescence micrographs of CEM cilia of wild-type and *rab-28(tm2636)* males expressing PKD-2::GFP. Dotted white boxes mark one CEM cilium. PKD-2 mainly localizes to the cell bodies, PCMC and cilium (ci) both of which are marked by white arrowheads. In *rab-28(tm2636)*, the PKD-2::GFP signal intensity is greater along anterior regions of the cilium. Note the difference in the distribution of the ciliary tip signal v/s the region posterior to it in *rab-28.* Scale bar is 5 μm for all panels in the figure. **(C)** Plot profile of PKD-2::GFP intensity across different points along the cilium in wild-type and *rab-28(tm2636)* adult males. Traces run from the ciliary tip and posterior towards the PCMC. Each data point represents the average PKD-2::GFP intensity at an individual point on several cilia in many animals of each genotype. *rab-28(tm2636)* males accumulate more PKD-2 anterior to the site where PKD-2 accumulation is greatest in wild-type. n=46 cilia and N=32 animals for wild-type, n=57 cilia and N=36 animals for *rab-28(tm2636).* **(D)** Graphs depicting the average PKD-2::GFP intensity across the whole cilium between WT and *rab-28(tm2636)* adult males. p=0.1640 calculated by Mann-Whitney test. n=41 and 50 cilia for wild-type and *rab-28(tm2636).* Data is from three separate trials. **(E)** Scatter plots depicting the number of PKD-2::GFP EVs surrounding the male head per animal between wild-type and *rab-28(tm2636).* p=0.2425 as calculated by Mann-Whitney test. N=47 and 48 animals for wild-type and *rab-28(tm2636)*. **(F)** Fluorescent micrographs of wild-type and *rab-28(tm2636)* males expressing CIL-7::GFP. Dotted white boxes mark one cilium. CIL-7 mainly localizes to the cell bodies, dendrites, PCMC and cilium(ci). PCMC and cilium are marked by white arrowheads. In *rab-28(tm2636)*, more CIL-7 is seen along/outside the cilium. **(G)** Plot profile of CIL-7::GFP intensity across different points along the cilium in wild-type and *rab-28(tm2636)* adult males. Traced similarly to 4F. *rab-28(tm2636)* males accumulate more CIL-7 along the length of the cilium compared to wild-type. n=48 cilia and N=34 animals for wild-type, n=32 cilia and N=18 animals for *rab-28(tm2636).* **(H)** Graphs depicting the average CIL-7::GFP intensity across the length of the cilium between WT and *rab-28(tm2636)* adult males. p<0.0001 calculated by a Mann-Whitney test. n=48 and 32 cilia for wild-type and *rab-28(tm2636).* Data is from three separate trials. **(I)** Scatter plots depicting the number of CIL-7::GFP EVs surrounding the male head per animal between wild-type and *rab-28(tm2636).* p=0.7728 by Mann-Whitney test. N=46 and 47 animals for wild-type and *rab-28(tm2636)*.

**Figure 5.**
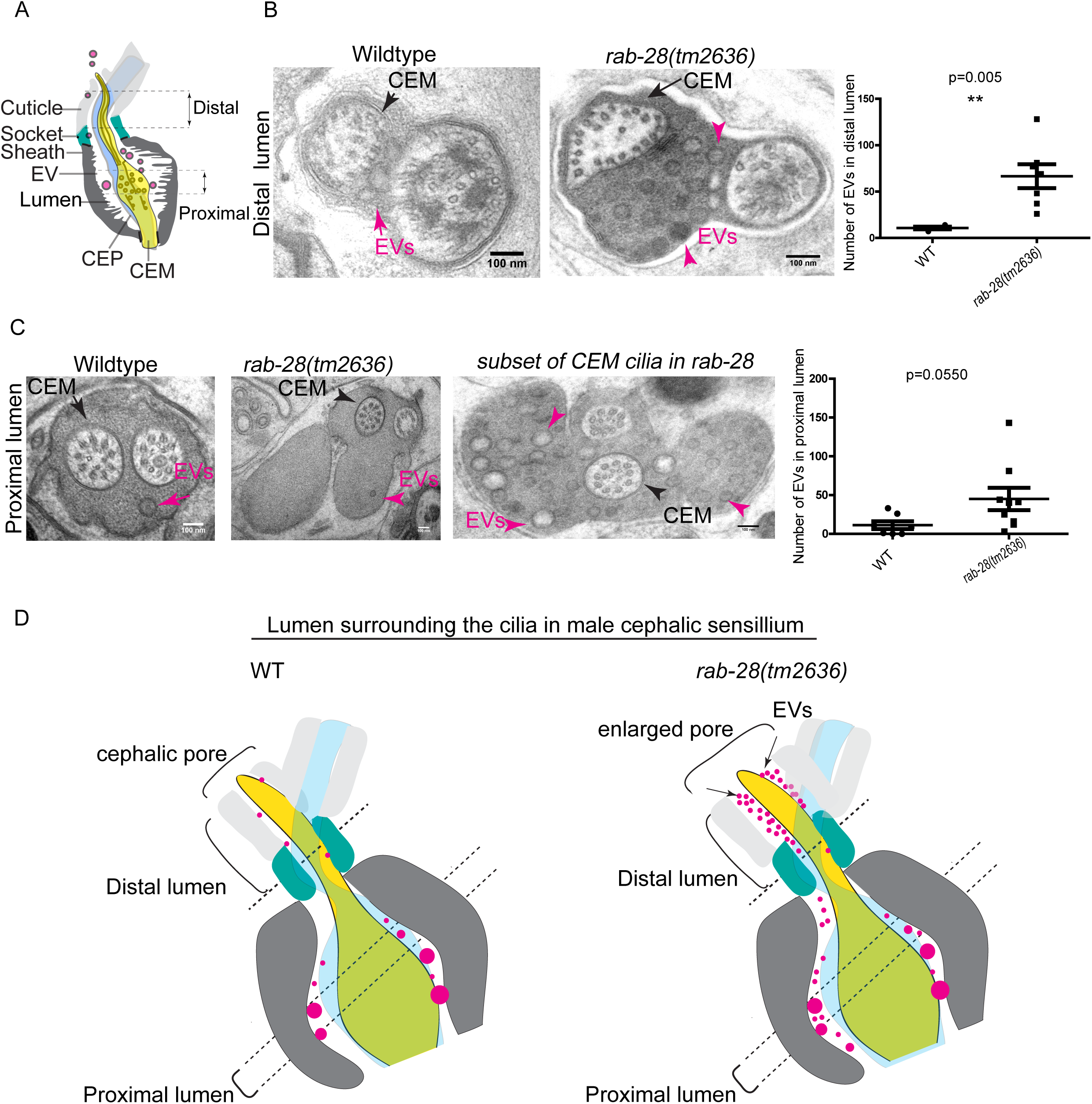
RAB-28 is a negative regulator of ciliary EV biogenesis and shedding. **(A)** Cartoon of the ultrastructure of the cephalic sensory organ reproduced from (J. Wang et al. 2014). EVs (magenta spheres) are ‘shed’ from the PCMC/ciliary base into the lumen and ‘released’ into the environment. **(B)** Transmission electron micrographs of the cephalic lumen surrounding the distal region of CEM cilia. Black arrows point to the CEM and magenta arrows to EVs. *rab-28(tm2636)* accumulate significantly more EVs in the lumen distal to the singlet region of CEM compared to wild-type males. p=0.005 as determined by unpaired t-test with Welch’s correction. n= 4 cilia and N= 2 animals for wild-type. n= 7 cilia and N= 3 animals for *rab-28(tm2636)*. Scale bars; 100nm **(C)** Transmission electron micrographs of the cephalic pore at the level of the CEM cilium transition zone. Scale bars; 100nm. A subset of the *rab-28(tm2636)* animals accumulate EVs in the cephalic lumen surrounding the TZ. p=0.0550 as calculated by unpaired t-test with Welch’s correction. The variances are significantly different as assessed by a F-test p =0.0098. **(D)** Cartoon depictions of the lumen surrounding the cilia in male cephalic sensillum in wild-type and *rab-28(tm2636)* sensilla. Color scheme is same as that of cartoon in Figure 5A. Brackets enclose cephalic sensory organ pore region. *rab-28* mutant males accumulate an excess of EVs (labeled by magenta spheres and pointed to by arrows) in the lumen surrounding the more distal portion of the CEM axoneme. *rab-28* mutants also have an enlarged cephalic pore.

PKD-2 is a TRP polycystin-2 that is expressed in male-specific EVNs, including the CEM neurons, and localizes to cilia and ciliary EVs (Barr et al. 2001; J. Wang et al. 2014). In wild-type males, PKD-2::GFP is observed at the PCMC and ciliary tip of CEM neurons, and within EVs in the cephalic lumen and those that are environmentally released (Figures 4B and 4C). In *rab-28(tm2636)* mutants, total PKD-2 levels at the CEM ciliary region from the PCMC to the tip are similar to wild-type (Figure 4D). However, fluorescence intensity measurement analyses revealed a subtle yet consistent change in the pattern of PKD-2::GFP distribution in *rab-28(tm2636)* worms indicating a ciliary localization defective (Cil) phenotype. Specifically, unlike wild-type animals where PKD-2::GFP signals are most intense at the PCMC level, *rab-28* mutants display elevated levels of PKD-2::GFP at more distal parts of the CEM cilium region, with reduced levels at the PCMC level (Figure 4B).

Cil defects could arise due to defects in PKD-2 trafficking to cilia or defects in PKD-2-positive EV biogenesis and release (J. Wang et al. 2014; Maguire et al. 2015; Silva et al. 2017; O’Hagan et al. 2017; Bae et al. 2006). Since the abnormal PKD-2::GFP accumulation at more distal regions of the *rab-28* cephalic pore appears to extend beyond that predicted by sole localization to the relatively narrow CEM cilium, we suspected that this excess GFP signal is due to abnormally high levels of PKD-2-positive EVs in the cephalic lumen. To determine if *rab-28* regulates environmental EV release similar to IFT and tubulin code regulators (J. Wang et al. 2014; Maguire et al. 2015; Silva et al. 2017; O’Hagan et al. 2017; Bae et al. 2006)), we compared the numbers of PKD-2::GFP-containing EVs in mounting media surrounding wild-type and *rab-28(tm2636)* males. We found that *rab-28(tm2636)* males release similar numbers of PKD-2::GFP-containing EVs as wild type, ruling out a function for RAB-28 in environmental EV release of PKD-2 and suggesting a possible defect in EV shedding (Figure 4E).

To further explore a role for *rab-28* in ciliary EV biogenesis and/or shedding, we examined the localization pattern of CIL-7, a peripheral membrane protein that localizes to cilia and ciliary EVs in all 27 EVNs (Maguire et al. 2015). In wild-type cephalic sensory organ, CIL-7::GFP localizes to the CEM neuronal dendrite, PCMC, cilium, and ciliary EVs. In contrast to the relatively subtle Cil phenotype with PKD-2::GFP, the *rab-28(tm2636)* cephalic sensory organ accumulates massive amounts of CIL-7::GFP in ciliary regions (Figures 4F-H). We observed a similar strong Cil phenotype in *rab-28(gk1040)* worms (Supplementary Figure 2). As is the case for PKD-2-containing EVs, wild-type and *rab-28* males release similar numbers of CIL-7::GFP-containing EVs in mounting media (Figure 4I and supplementary Figure 2C). Together, our results indicate a role for *rab-28* in regulating the abundance and distribution of ciliary proteins expressed in the EV-releasing CEM neurons. Furthermore, our results show that *rab-28* is not required for the release of PKD-2- and CIL-7-positive EVs into the environment.

### RAB-28 negatively regulates EV shedding in cephalic sensory organs

To determine if the *rab-28* Cil phenotype is caused by defects in CEM ciliary axoneme or with EV biogenesis and shedding, we examined the ultrastructure of the four cephalic sensory organs in males cryofixed via high pressure freezing (HPF). In the cephalic male sensillum, the cilia of the CEM neuron and its neighboring CEP neuron are enclosed in a pore formed by the glial cephalic sheath and socket cells and overlying cuticle (Figure 5A). The CEM ciliary tip is exposed to the external environment through the pore that contains a cuticular plug. In TEM images of wild-type cephalic male sensory organs, EVs are observed in the lumen, which is an extracellular space within the pore (Figures 5B and 5C, Supplemental Movie 3). The CEM cilium at the distalmost segment is comprised only of singlet microtubules and pokes through the cuticle to access the environment. Posterior to this, the CEM and the CEP cilia are jointly surrounded by the cuticle. In wild-type males, the cephalic lumen at this level contains very few EVs, and those EVs are usually clear and often irregular in shape and size (Figure 5B) (J. Wang et al. 2014; Silva et al. 2017). In *rab-28(tm2636),* the distal pore is enlarged and the cephalic lumen accumulates an excess of EVs that are smaller and more uniform in size and sometimes darkly stained possibly indicating a change in composition (Figure 5B, Supplemental Movie 3). Posterior to this, the axoneme of the CEM cilia is surrounded by the socket glia, and is comprised mainly of A-tubule and B-tubule singlets that are formed by the characteristic split of AB doublet microtubules — a feature that may be crucial to EV release (Silva et al. 2017; O’Hagan et al. 2017). Consistent with a lack of an EV release defect in *rab-28* mutants, we did not observe any defects in the specialized CEM doublet microtubule split.

To determine whether the excessive and ectopic EV accumulation in *rab-28* mutant males was due to an occlusion of the pore or a ciliary tip defect, we examined the distal segments of *rab-28* CEM cilia using electron tomography. Similar to wild-type, we found that *rab-28* cilia are indeed exposed to the environment (Supplementary Figure 3). We interpret the lack of a defect in the ciliary tip, cephalic pore environmental access or environmental EV release to mean that the EV accumulation phenotypes in *rab-28* worms are not due to blockage of shed but unreleased EVs.

The sheath glial cell surrounds most of the doublet region, TZ, and PCMC of the CEM cilium. Similar to excessive EVs surrounding distal cilia, a subset of *rab-28* cephalic sensilla accumulate EVs that are greater in number than the maximum number of EVs ever observed in wild-type (Figure 5C). Some of these EVs are darker and uniform in size in contrast to wild-type EVs, which are clear and heterogeneous in size. We did not find the increase in EV numbers surrounding *rab-28(tm2636)* TZ and PCMC to be statistically significant (Figure 5C, Supplemental Movie 3). We also did not observe any striking axonemal ultrastructural defects at the ciliary doublet or TZ levels in *rab-28(tm2636)* worms. Given the conserved localization of RAB-28 to the PCM of cilia, we also compared the PCMC of wild-type and *rab-28(tm2636)* males. We observed no gross ultrastructural differences, although a higher incidence of darker EVs in the lumen surrounding the *rab-28* PCMC was noted, consistent with the appearance of EVs surrounding other regions of CEM axoneme. We also observed a noticeable but not significantly different increase in the number of EVs in the lumen surrounding the PCMC (Supplementary Figure 4).

Combining our live imaging and ultrastructural analyses, we conclude that *rab-28* mutant males produce and shed excessive amounts of abnormally stained and sized EVs into the cephalic sensory organ. These data suggest that RAB-28 regulates EV cargo sorting and production (biogenesis) and that RAB-28 also acts as a negative regulator of EV shedding without affecting environmental EV release.

### BBSome and *arl-6*, but not PDL-1, loss phenocopies rab-28 mutant EV shedding defects

To further explore EV-related roles and mechanism of RAB-28 action, we assessed ciliary EVs in RAB-28 transport regulator mutants. Similar to *rab-28* mutants, we found Cil defects in *bbs-8* and, to a lesser extent, *arl-6* mutants, with the ciliary EV marker CIL-7::GFP accumulating at the PCMC and around the axonemal region of CEM cilia (Figure 6A-C). This observation is consistent with a similar, previously reported, PKD-2::GFP Cil phenotype in *bbs-7* mutants (Braunreiter et al. 2014). In contrast, *pdl-1* mutants show no gross CIL-7 localization defects (Figure 6A and quantified in 6C). We also examined whether *bbs-8, arl-6* or *pdl-1* regulate the release of CIL-7::GFP-marked EVs. Similar to *rab-28* mutant males, we found no differences in the number of CIL-7-positive EVs released by *bbs-8, arl-6* and *pdl-1* mutants when compared to wild-type males (Figures 6D and 6E).

**Figure 6.**
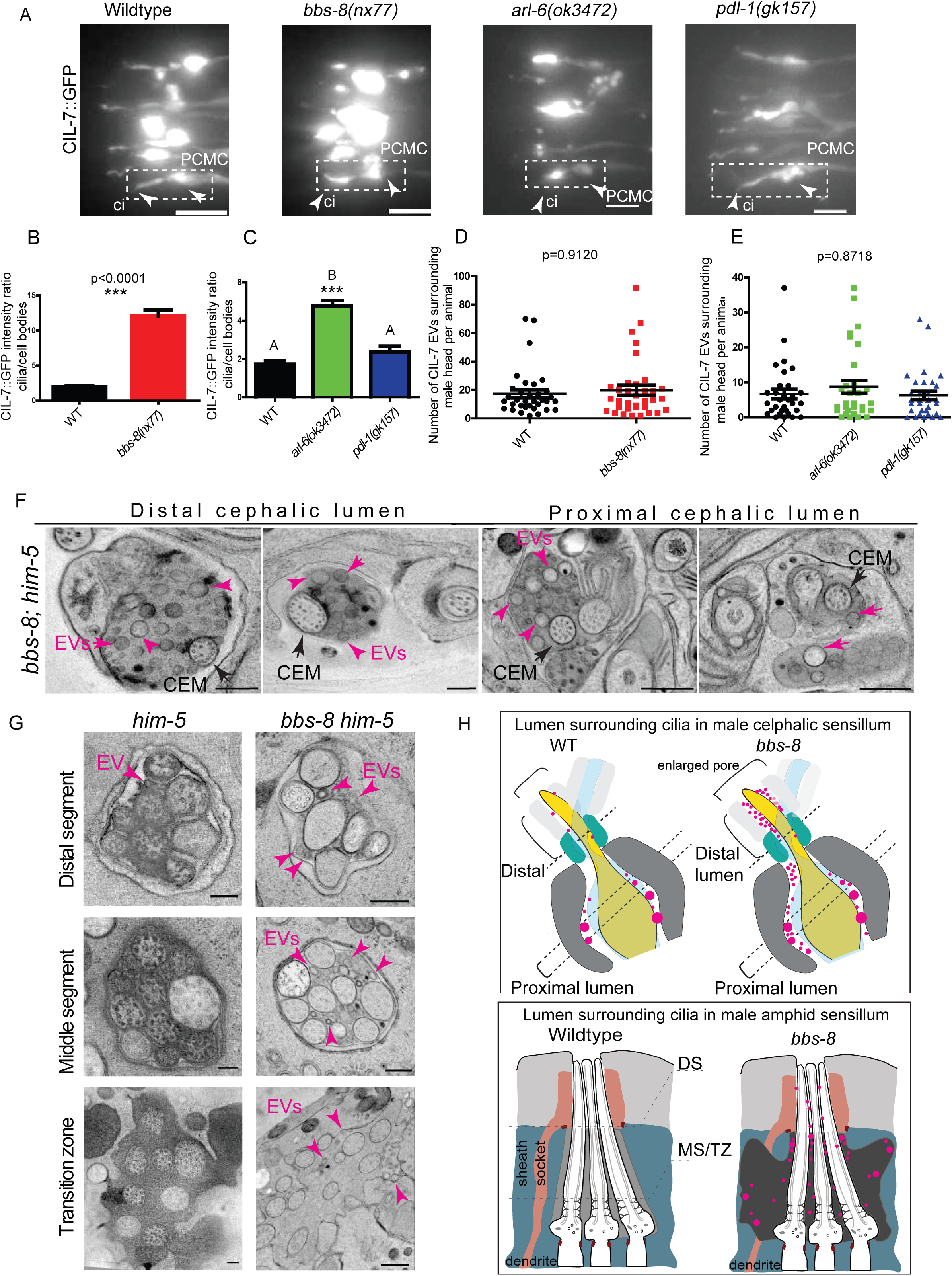
Localization of RAB-28 to the PCM is essential to regulate EV shedding. **(A)** Fluorescence images of CIL-7::GFP in the male heads of wild-type, *bbs-8(nx77), arl-6(ok3472)* and *pdl-1(gk157)* mutants. Scale bars; 5 μm. **(B)** Graph depicting the ratio of CIL-7::GFP intensity between the ciliary and cell body regions in wild-type and *bbs-8(nx77)* mutants. p<0.0001 as determined by Mann-Whitney test. N=21 animals for both genotypes. **(C)** Graph depicting the ratio of CIL-7::GFP intensity between the ciliary and cell body regions in wild-type, *arl-6(ok3472)* and *pdl-1(gk157)* males. Letters above each group indicate results of statistical analysis; data sets that do not share a common letter are significantly different at p<0.005 based on Kruskal–Wallis test with Dunn’s post-hoc correction. *arl-6* mutants accumulate more CIL-7::GFP in heads. N=27 animals for all genotypes **(D)** Scatter plot depicting the number of CIL-7::GFP-labeled EVs released from wild-type and *bbs-8* heads. p=0.9120 as determined by Mann-Whitney test. N=34 animals for both genotypes. **(E)** Scatter plot depicting the number of CIL-7::GFP EVs released from wild-type, *arl-6(ok3472)* and *pdl-1(gk157)* heads. p=0.8718 as determined by Kruskal-Wallis test with Dunn’s multiple comparisons. N=31 animals for all genotypes. **(F)** TEM cross sections of cephalic lumen of *bbs-8* mutant males.Black arrows point to CEM. Ectopic EVs (magenta arrows) are observed at distal and proximal regions of the lumen. Scale bar; 200nm. **(G)** TEM cross sections of the lumen surrounding amphid channel cilia in *bbs-8* mutant males, showing ectopic accumulated EVs (magenta arrows) at distal and middle segments and around the transition zones. Scale bar; 200nm. **(H)** Cartoon depictions of cephalic sensilla in wild-type and *bbs-8* mutants. *bbs-8* mutants accumulate an excess of EVs (magenta spheres) in the lumen surrounding all levels of the CEM axoneme. Brackets on top mark cephalic pore size. *bbs-8* mutants have enlarged cephalic pores. Bottom cartoons depict amphid pore ultrastructure phenotypes in wild-type and *bbs-8* mutants. *bbs-8* amphid cilia in male *C.elegans* accumulate EVs (magenta spheres) during adulthood.

To directly determine whether the BBSome regulates EV shedding, we examined the ultrastructure of the cephalic sensory organs of cryofixed (HPF) *bbs-8* mutant males. As in *rab-28* mutants, we found that the *bbs-8* cephalic lumen was distended and filled with EVs, both at the distal end and at the level of the ciliary transition zone and PCMC (Figure 6F). *bbs-8* mutants accumulate more EVs than *rab-28*, and the accumulated EVs are heterogeneous in size in contrast to the uniformly sized EVs found in the distal lumen of *rab-28* mutant males. *bbs-8* mutant CEM cilia have no gross axonemal defects at the TZ similar to *rab-28*. While our TEM analysis of sub-distal CEM axonemes in *bbs-8* mutants indicated that the conserved AB tubule split is unaffected, we were limited by the availability of complete serial sections covering that region thus rendering our analysis of this ciliary segment incomplete. We also observed ECM deposits within a subset of cephalic sheath glia, matching the amphid sheath cell phenotype of *bbs-8* mutants. Strikingly, the amphid pores of cryofixed *bbs-8* mutant males were also filled with EVs (Figure 6G), which were not detectable in chemically fixed hermaphrodite samples (Figure 2B, C). These EVs were observed at all levels of the *bbs-8* male amphid pore, including the distal and middle segment, TZ, and PCMC regions. Thus, like RAB-28, the BBSome also negatively regulates EV shedding, although this role appears to extend beyond the EVNs to include the amphid neurons.

Together, these data indicate that BBS-8 and ARL-6, but not PDL-1, regulate the ciliary localization and levels of the EV marker, CIL-7. We also confirm that the *bbs-8* CIL-7 defect correlates with an EV shedding defect, indicating that the BBSome and RAB-28 share a common role in inhibiting EV shedding. Notably, the PCM localization phenotype for RAB-28^Q95L^ correlates with the Cil and EV phenotypes in mutants of *bbs-8* (no PCM localization; strong Cil and EV defects), *arl-6* (reduced PCM localization; partial Cil defect) and *pdl-1* (normal PCM localization; no Cil defect). Combined, these results suggest that the main site at which RAB-28 acts to regulate EV shedding is likely to be the PCM.

## DISCUSSION

### An integrated BBSome, ARL-6 and LIFT pathway regulates the ciliary targeting of RAB-28

We present here a network of ciliary trafficking pathways that cooperate to regulate RAB-28 levels both at the periciliary membrane and within cilia. We show that BBS-5 promotes RAB-28 PCM and IFT association, thereby solidifying the requirement of the BBSome for regulating RAB-28 localization originally described (Jensen et al. 2016). We find that the orthologue of the mammalian BBSome regulator, ARL-6, also maintains normal RAB-28 PCM levels, albeit not to the same extent as the BBSome. Interestingly, *arl-6* mutants show an abnormally high frequency of RAB-28-positive IFT trains in amphid and phasmid channel cilia, which could explain why steady state levels of RAB-28 at the PCM are reduced in this strain. Thus, in contrast to BBSome components, ARL-6 negatively regulates RAB-28 association with IFT trains. In addition, we discovered that RAB-28 association with IFT, but not the PCM, requires PDL-1 and its presumptive cargo displacement factor ARL-3. Notably, *bbs-8* and *arl-6* are epistatic to *pdl-1* for RAB-28 PCM localization, suggesting that the BBSome functions upstream of PDL-1 in the ciliary RAB-28 trafficking pathway. Based on these data, we propose a model whereby the BBSome targets RAB-28 to the PCM, after which PDL-1 solubilizes the G-protein as a prerequisite for association with IFT trains, a step that is negatively regulated by ARL-6.

Intriguingly, we observe cell-type specific differences in this pathway. In male EVN cilia, the frequency of RAB-28^Q95L^ IFT events is on average one fifth of that in phasmids. Whilst the requirement for the BBSome in regulating RAB-28 PCM association and IFT behavior is identical in all analyzed ciliary subtypes, we find that PDL-1 is partially required for maintaining RAB-28 PCM levels in EVN cilia but not amphid/phasmid cilia. In addition, a synthetic relationship is observed for *pdl-1* and *arl-6*, in that RAB-28 is fully delocalized from the PCM region of *pdl-1;arl-6* double mutant EVNs. The reasons for such distinctions in the deployment of RAB-28 trafficking machinery in different cell types are unclear, although it may be due to differences in how RAB-28 functions in CEM cilia versus amphid/phasmid cilia and due to the fact that an additional pool of RAB-28 is maintained in the distal regions of CEM cilia but not amphid cilia.

Thus, we have uncovered a network consisting of the BBSome, ARL-6 and components of the LIFT system that regulate RAB-28’s PCM and ciliary localization, and IFT association, in a cell-specific manner. RAB-28 is one of only two proteins known to undergo both IFT and LIFT, the other being INPP5E (Fansa et al. 2016; Kösling et al. 2018). In both cases, LIFT is required for initial ciliary import and IFT for subsequent transport within cilia and, presumably, exit. This may, therefore, be a general mechanism for the ciliary transport of lipidated proteins. Aside from its functional roles in cilia, RAB-28 could serve as a useful model for investigating the trafficking of ciliary lipidated proteins and crosstalk between IFT and LIFT.

### RAB-28 and the BBSome negatively regulate EV shedding *in vivo*

We previously proposed that RAB-28 ciliary function could be via a role in EV biology (Jensen et al. 2016). Here, we show that RAB-28 regulates EV biogenesis and shedding in cephalic sensory organs, which display massive ectopic EV accumulation in *rab-28* mutants. We also show that defective targeting of RAB-28 to the PCM, via mutation of the BBSome, elicits a similar phenotype. This finding indicates: (i) RAB-28 and the BBSome serve as a negative regulator of EV shedding, and (ii) the PCM may be the site of RAB-28’s EV-related function. Interestingly, we find a CEM cilia specific dependence on IFT-association of RAB-28 in its enrichment at the PCM, consistent with a role for IFT in the regulation of EV shedding (J. Wang et al. 2014). Our findings indicate a tuneable mechanism consisting of transport regulators that control the levels and localization of EV regulators in cells and cilia.

Whilst the EV phenotypes of *rab-28* and *bbs-8* mutants clearly overlap, there are also some differences in terms of the pattern and severity. In *rab-28* males, EVs predominantly accumulate around more distal parts of the CEM cilium, whereas *bbs-8* males display a more severe phenotype with excess and more heterogeneous EVs surrounding *all* parts of the CEM cilium. Furthermore, EV accumulation occurs in the amphid pore of *bbs-8*, but not *rab-28*, mutant males. These differences suggest that the BBSome regulates the biogenesis of EVs beyond those that are dependent on RAB-28, and indicates the presence of additional EV regulators in CEM cilia. Future studies will exploit the diversity of CEM ciliary EV cargoes to test the cargo specificity of BBSome and RAB-28 in EV biogenesis.

Although the mechanism of RAB-28-mediated EV regulation remains unknown, our data indicate a connection between RAB-28 and the BBSome as negative regulators of EV shedding. Consistent with this functional relationship, RAB-28 in *Trypanosoma brucei* plays a role in the targeting of cargoes to lysosomes for degradation, a function also attributed to the BBSome in *C. elegans* (Lumb et al. 2011; Xu et al. 2015). In cultured mammalian cells, defective retrieval of sensory receptors due to loss of BBSome function leads to those cargoes being directed to ciliary-derived EVs called ectosomes (Nager et al. 2017). Thus, it is possible that excessive EV shedding in *bbs-8* and *rab-28* mutants could arise due to changes in endocytic events at the PCMC.

Interestingly, RAB-28 shares sequence similarities with IFT27 that plays a role in the exit of BBSome from cilia (Liew et al. 2014; Eguether et al. 2014; Jensen et al. 2016). Loss of IFT27 leads to EV shedding phenotypes due to its effects on the BBSome (Nager et al. 2017). One intriguing hypothesis is that *rab-28* EV phenotypes are indicative of BBSome dysfunction in CEM cilia. Since *C. elegans* lacks an IFT27 orthologue, RAB-28 could be important for BBSome functions in more distal parts of the CEM cilia, which corresponds spatially to that portion of the cephalic pore where EV phenotypes are most prevalent in *rab-28* mutants. While our transport data clearly supports the model that RAB-28 is downstream of the BBSome for its PCM and ciliary localization in amphid and CEM neurons, our EV shedding data raises a possibility that in certain ciliary contexts, RAB-28 might function similar to IFT27 and affect BBSome functions. However, whether *rab-28* and *bbs-8* regulate EV shedding using a similar mechanism remains to be determined.

### A role for EVs in ciliated neuron-glia communication

We observed defects in the pore of cephalic and amphid sensory organs, formed by the sheath and socket glia, in ectopic EV shedding mutants *rab-28* and *bbs-8*. These defects include pore volume expansion and large deposits of dense vesicular material in the sheath cytoplasm. Such deposits have been previously observed in the amphid sheath cells of ciliary mutants (Perkins et al. 1986; Barr et al. n.d.) and are thought to correspond to sheath-synthesized matrix-filled vesicles that deliver ECM to the amphid pore (Ward et al. 1975; Bacaj et al. 2008). A balance between exocytosis and endocytosis by the sheath cell regulates the amount of matrix deposited, which in turn determines pore volume (Perens & Shaham 2005). Signaling from cilia has been proposed to regulate the localization of glial factors such as DAF-6 and LIT-1 that regulate pore/sensory compartment size in amphid neurons (Oikonomou et al. 2011). Our data are consistent with a change in neuron-glia interactions in EV shedding mutants. We propose that ciliary signaling via RAB-28 and the BBSome coordinates morphogenesis of pores in sensory organs of *C. elegans*. Although the nature of the neuronal signal that participates in pore volume regulation and stimulates changes in the secretion of matrix from glia has not been identified, our previous work on the *rab-28*-disrupted amphid pore proposed it to be in the form of EVs (Jensen et al. 2016). EVs are implicated in many facets of nervous system development, signaling, and function (Zappulli et al. 2016; Budnik et al. 2016). EV exchange between neurons and glia has been documented *in vitro* and *in vivo* (Lopez-Verrilli et al. 2013; Frühbeis et al. 2013; Fröhlich et al. 2014; Goncalves et al. 2015). *C. elegans mec-9* ECM mutant males accumulate excess EVs with pore size defects similar to *bbs-8* supporting a possible relationship between EVs and pore size regulation (Barr et al. n.d.). Interestingly, the actin cytoskeleton that plays a role in pore/compartment size regulation in *C. elegans* is also important for EV release from cultured mammalian cilia (Oikonomou et al. 2011; Phua et al. 2017; Nager et al. 2017). We interpret the EV shedding defects in mutants that have enlarged pores and excess matrix to mean one or all of the following: under normal conditions, ciliated neurons release EVs that signal to the sheath glia to regulate pore size and matrix deposition; under normal conditions, ciliated neurons and glia communicate via EVs; in pathological conditions, excessive EVs may reflect abnormal neuron-glia communication consistent with altered pore size and excess matrix. It is important to note that whilst EVs are only very rarely observed in WT amphid pores, which is why the associated neurons are not considered EVNs, this pore clearly possesses the capacity to produce or take up EVs under certain pathogenic conditions (e.g, BBSome loss), in a sex-specific manner, or at specific developmental time-points. Indeed, our analyses have been limited to the adult male, leaving open the intriguing question of whether EVs play a role in the early development, morphogenesis, homeostasis, and/or function of sensory organs.

### EV shedding regulators in photoreceptor outer segment turnover and retinal degeneration

*rab28* knockout mice exhibit elongated cone outer segments (Ying et al. 2018). The discs of photoreceptor outer segments are themselves modified ciliary ectosomes (Salinas et al. 2017), raising the possibility that outer segment elongation could be a result of excessive disc formation in *rab28* mutant cones. Tantalizing as this hypothesis is, however, it does not explain the reduction in disc shedding from the outer segment in *rab28* knockout mice. An alternative explanation for the elongated outer segment could be that mammalian Rab28 has no role in disc formation, but solely regulates shedding, making it a positive regulator of this process. Whether human *RAB28*-associated cone-rod dystrophy is a disease of ciliary EV shedding remains to be seen. The apparent absence of extraocular phenotypes in both mice and humans deficient for RAB28 suggests either: (i) the retina is more sensitive to perturbation of ciliary EV shedding than other tissues (as it is for other ciliary functions), or (ii) RAB28 alone is not required for EV shedding outside the retina, consistent with phenotypic differences we observe in *C. elegans rab-28* mutant EV phenotype in different sensory organs (cephalic vs amphid).

The role of the BBSome in disc shedding, if any, is not known, as the BBSome is also essential for outer segment morphogenesis and maintenance (Hsu et al. 2017), precluding the analysis of shedding in BBSome mutants. EVs have been observed surrounding rod cilia in *bbs8* mutant mice (Dilan et al. 2018), a phenotype also observed in mutants of the EV release-suppressing Peripherin-2 (Salinas et al. 2017). Intriguingly, much like our hypothesis of pore size regulation in *C. elegans*, disc shedding is an example of the exchange of ciliary EVs between two cells. Thus, the EV shedding cilia of *C. elegans* could provide a simple model for the identification of molecules important for outer segment disc formation and turnover.

### Conclusions and open questions

Our work adds RAB-28 to a growing list of G-proteins involved in EV biogenesis and release that includes ARF6, RAB-11, RAB22A, RAB27 and RAB35 (Muralidharan-Chari et al. 2009; Wehman et al. 2011; Ostrowski et al. 2010; Hsu et al. 2010; T. Wang et al. 2014). An important question that remains unanswered is whether the EVs shed in *C.elegans* cephalic lumen and environmentally released are exosomes, ectosomes, or both. Genetic evidence coupled with the absence of MVBs in the PCM and cilium is consistent with ectosomal identity (J. Wang et al. 2014). While the majority of Rabs associated with EV biogenesis function in exosome formation and trafficking (Blanc & Vidal 2018), ciliary EVs are generally thought to be ectosomes (Wood et al. 2013; Nager et al. 2017). RAB-28 is only the second Rab family member shown to have a role in ciliary EV biogenesis, the other being IFT27/RABL4 (Nager et al. 2017). Neither is associated with EV generation in other contexts. Therefore, the cilium may use unique EV-regulators, enabling the cilium to modulate its EV output independently of the rest of the plasma membrane.

Our studies raise several interesting questions— How and why is PCM localized RAB-28 important to suppress EV shedding surrounding *distal* parts of cilia? What is the mechanism by which RAB-28 and the BBSome regulate EV shedding? Why aren’t RAB-28 and the BBSome packaged into EVs despite being closely associated with the biogenesis process? How are proteins sorted and packaged into ciliary EVs? What are the effectors and GAP/GEF of RAB-28? How do ciliary EVs interact with target cells such as glia? Answers to these questions will lead to a better understanding of the fundamental biology of ciliary EVs, neuron-glia signaling, and ciliopathies with relatively unexplored EV phenotypes such as polycystic kidney disease, Bardet-Biedl Syndrome, and autosomal recessive cone-rod dystrophy.

## AUTHOR CONTRIBUTIONS

JA, SC, MMB and OEB conceived the project, designed the experiments and interpreted the results. JA conducted and analyzed ciliary EV cargo localization and EV release in all mutants. FR constructed EV cargo reporter strains. JA, KCQN and DDH did cryofixations on *rab-28* and *bbs-8* mutant males, and performed TEM on *rab-28* mutants. JA and MS generated 3D models of cephalic sensilla. SC and ALM conducted and analyzed RAB-28 localization experiments. SC analyzed all BBSome and *arl-6* mutant TEM. ST performed TEM on *bbs-8* mutants. JA and SC compiled the figures and wrote a first draft of the manuscript. JA, SC, MMB and OEB wrote the final manuscript. MMB, BNK and OEB supervised the research.

## ACKNOWLEDGEMENTS

This work was funded by the National Institutes of Health (NIH) awards DK059418 and DK116606 to MMB and OD010943 to DHH, Science Foundation Ireland (SFI) principal investigator (11/PI/1037) and SFI-BBSRC (Biotechnology and Biological Sciences Research Council) partnership (16/BBSRC/3394) awards to OEB, and an Irish Research Council (IRC) Government of Ireland postgraduate award (GOIPG/2014/683) to SC, a New Jersey Commission on Spinal cord research (NJCSCR) postdoctoral fellowship (CSCR16FEL008) to JA. We thank Juan Wang for discussion and insight on ciliary EVs, Gloria Androwski and Helen Ushakov for excellent technical assistance, Barr labmates and the Rutgers *C. elegans* community for feedback and constructive criticism throughout this project, the Conway Institute imaging core facility for technical support, and WormBase. We thank Leslie Gunther-Cummins and Xheni Nishku at AECOM for assistance with high pressure freeze fixation. We also thank the National BioResource Project (Tokyo Women’s Medical College, Tokyo, Japan) and *Caenorhabditis* Genetics Center (CGC) for strains. The CGC is supported by the National Institutes of Health - Office of Research Infrastructure Programs (P40 OD010440).

---

**Supplementary Figure 1.**
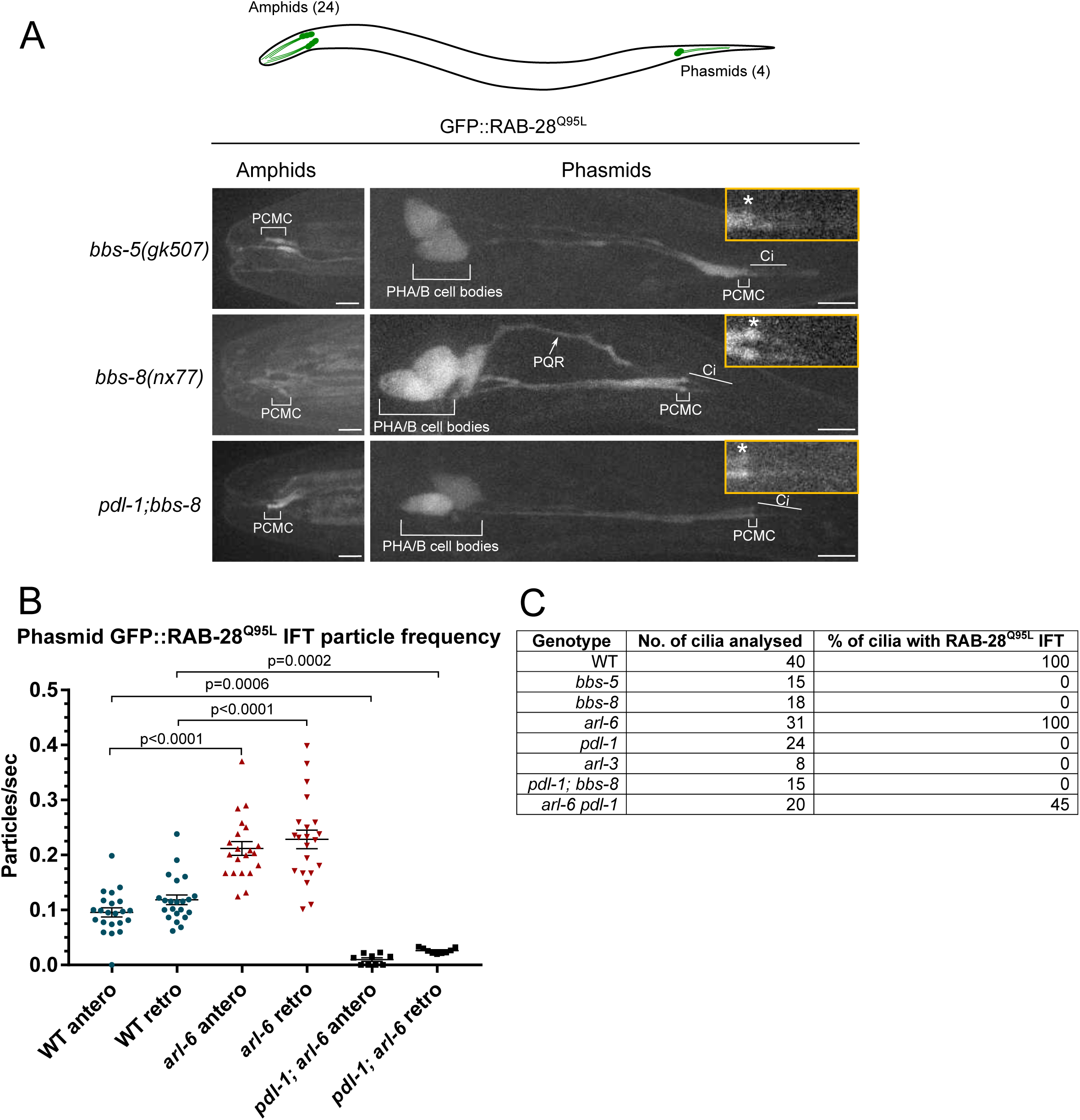
GFP::RAB-28 localization in BBSome mutants, *arl-6* and *pdl-1* regulate the frequency of RAB-28 IFT events. **(A)** Representative confocal z-projection images of amphid and phasmid neurons from worms expressing GFP::RAB-28^Q95L^. Images are the same as those in Figure 1 except that the intensity has been increased to visualize the mislocalized GFP signals. Insets show blow ups of phasmid cilia, asterisks indicate PCMC. Anterior is to the left. Scale bars; 10 μm. **(B)** Scatter plots showing the frequency of anterograde and retrograde GFP::RAB-28 IFT particles, in particles per second, as derived from kymographs taken from N2 (WT), *arl-6* and *pdl-1; arl-6* phasmid cilia expressing GFP::RAB-28^Q95L^. The frequency with which GFP-RAB28^Q95L^ undergoes IFT is effectively doubled in the *arl-6* background, but reduced to 7.8% (anterograde) and 16.4% (retrograde) of WT in the *pdl-1; arl-6* background, on average. Error bars show SEM; p-values were calculated by one-way ANOVA with Tukey’s multiple comparison test. Graphed data are from 22 N2, 21 *arl-6* and 9 *pdl-1; arl-6* kymographs. **(C)** Table summarizing the percentage of phasmid cilia with detectable IFT movement of GFP::RAB-28^Q95L^ in the indicated mutants.

**Supplementary Figure 2.**
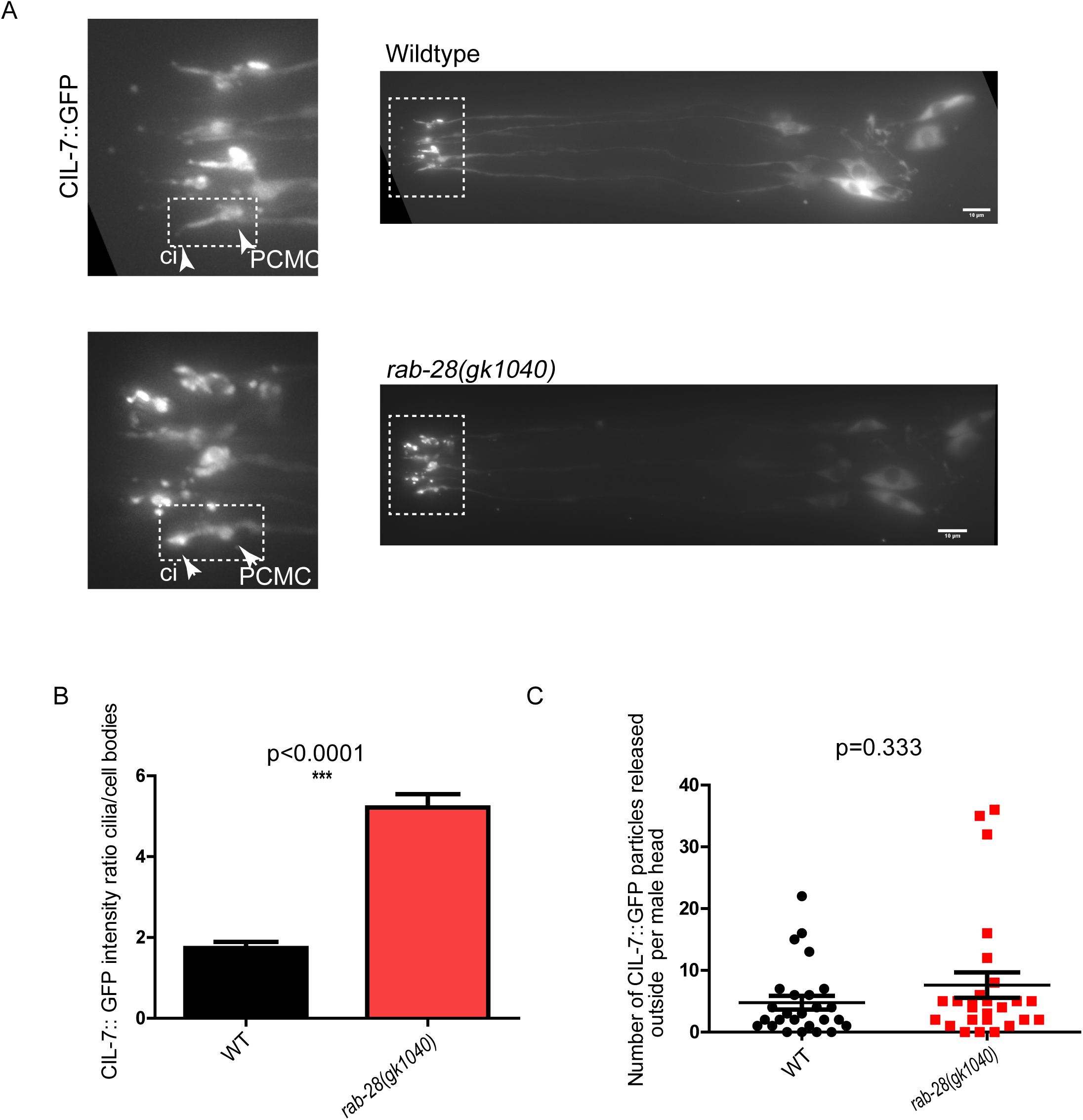
*rab-28(gk1040)* phenocopies the Cil phenotype of *rab-28(tm2636)* worms. **(A)** Fluorescence micrographs of wild-type and *rab-28(gk1040)* adult males expressing CIL-7::GFP. Close-ups of the male heads are shown on the left. Dotted white boxes mark one cilium. White arrowheads in insets mark PCMC and cilium (ci). In *rab-28(gk1040)*, more CIL-7 is seen along the cilium. Scale bar; 10μm. **(B)** Graph depicting the ratio of CIL-7::GFP intensity between cilia and cell bodies in wild-type and *rab-28(gk1040).* Similar to *rab-28(tm2636)*, *rab-28(gk1040)* males also accumulate CIL-7::GFP along the cilium. p<0.0001 as determined by Mann-Whitney test. N=27 animals for both genotypes. **(C)** Scatter plots depicting the number of CIL-7::GFP EVs surrounding the male head per animal between wild-type and *rab-28(gk1040).* p=0.333 for CIL-7 EV comparisons by Mann-Whitney test. N=31 animals for both genotypes.

**Supplementary figure 3.**
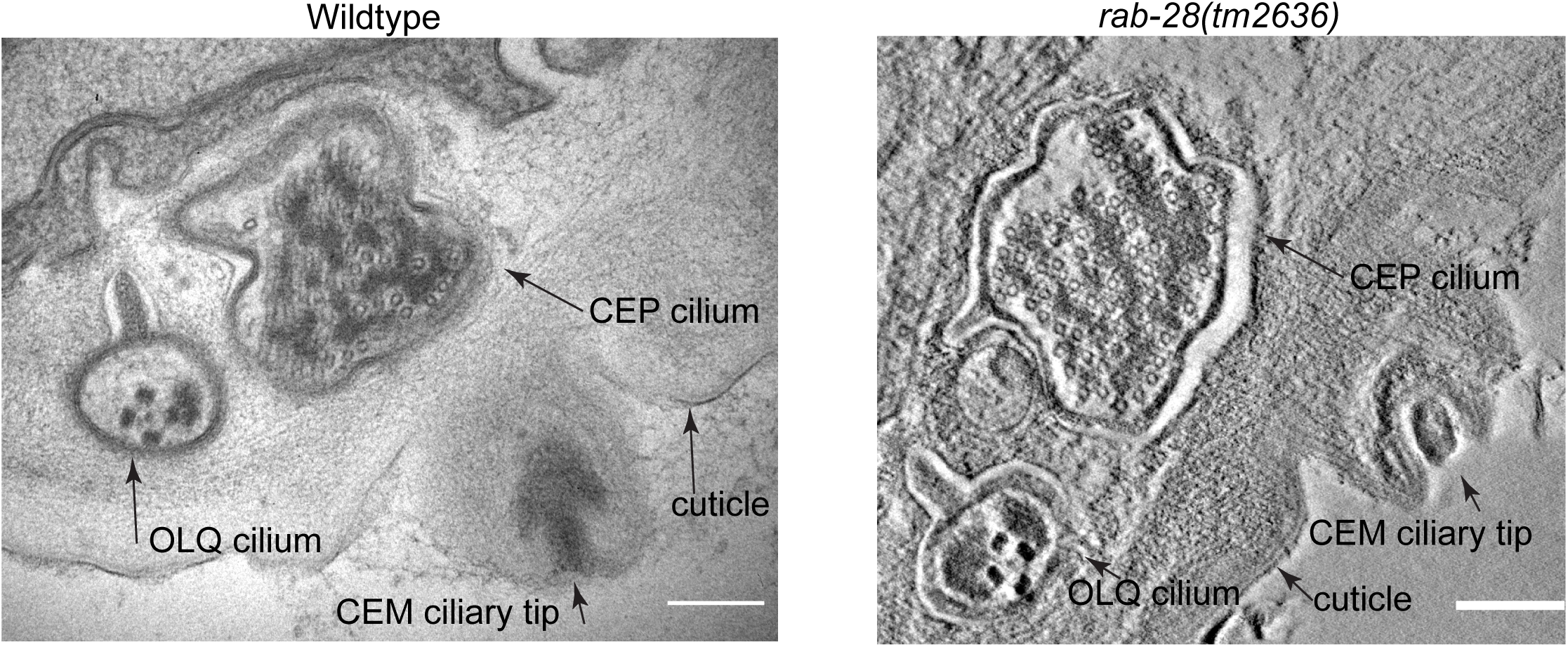
TEM of wild-type (*him-5)* CEM ciliary tip (left) and slice view of an electron tomogram of *rab-28(tm2636)* CEM ciliary tip (right). Arrows point to the CEM ciliary tips, its neighboring CEP cilium, the cuticle overlying the CEM cilium, and the nearby OLQ cilium. Note that the CEM ciliary tip is open and exposed to the environment in both genotypes. Scale bar is 200nm.

**Supplementary figure 4.**
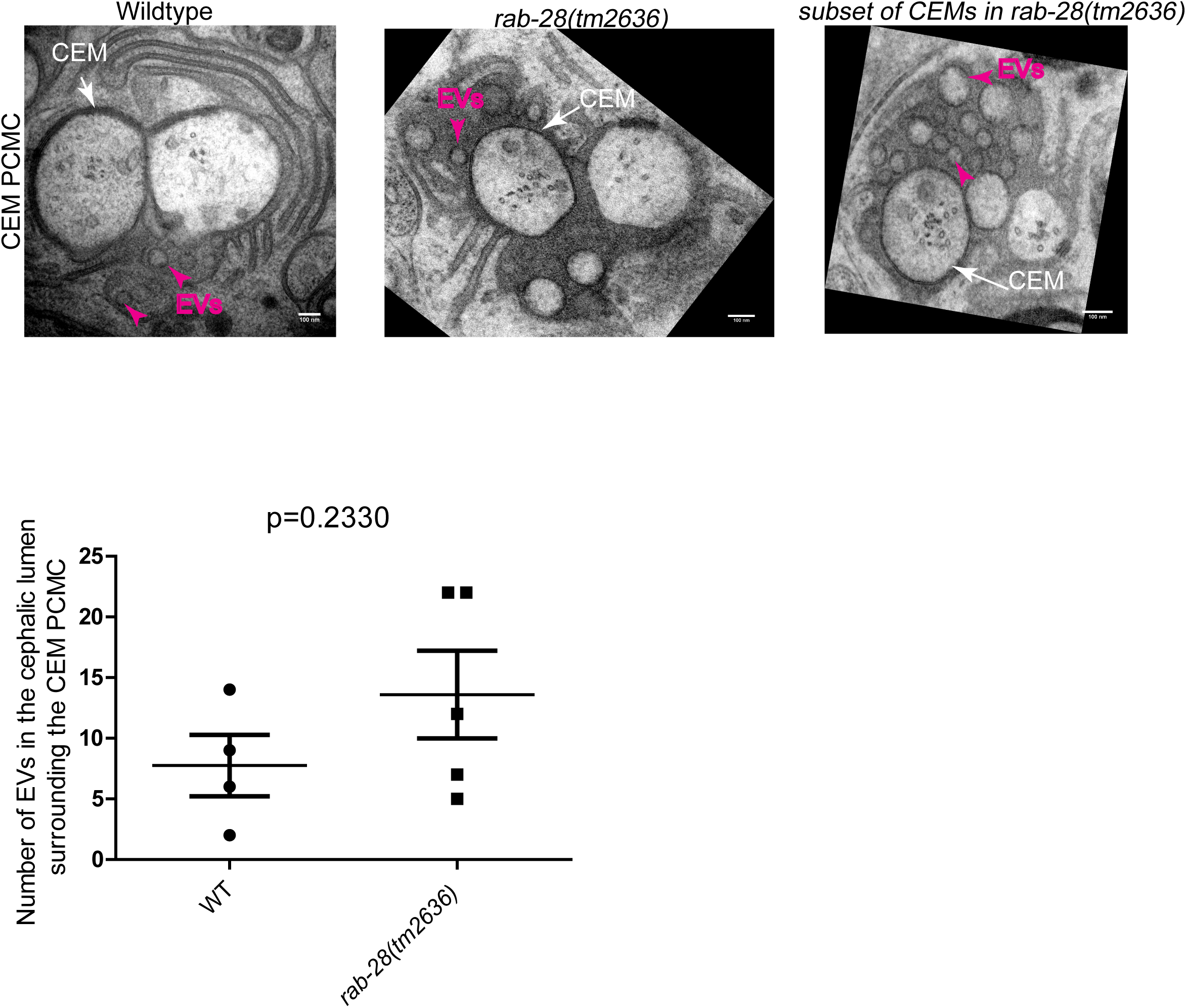
TEM of wild-type (*him-5)* and *rab-28(tm2636); him-5* cephalic lumens at the PCMC level. White arrows point to the CEM neurons. A subset of *rab-28* accumulate more EVs (magenta arrowheads) in the cephalic lumen. Scale bar is 100nm. Bottom-scatter plot comparing the number of EVs accumulating in the cephalic lumen at the PCMC level. p=0.2330 as analyzed by an unpaired t-test with Welch’s correction. n=4 cilia and N=2 animals for wildtype, n=5 cilia and N=2 animals for *rab-28(tm2636)*.

**Supplementary movie 1.** Representative movies of GFP::RAB-28^Q95L^ IFT behavior in the phasmid cilia of N2 (WT), *arl-6* and *pdl-1; arl-6* mutants. Anterior is to the left. Movies are played at 5 fps. Scale bars; 2 μm. PCMC: periciliary membrane compartment.

**Supplementary movie 2.** Representative movies of GFP-RAB-28^Q95L^ IFT behavior in CEM and RnB cilia of *him-5* males. A RAB-28-positive IFT particle can be seen in the bottom CEM and RnB cilia. Higher frequency IFT movement of RAB-28 is evident from the amphid cilia. PCMCs are marked by asterisks. Movies are played at 6 fps. Anterior is to the left. Scale bars 2 μm.

**Supplementary movie 3.** Electron tomography and serial section TEM based model of the male cephalic sensillum of wild-type and *rab-28(tm2636)* respectively. Models depict the CEM cilium (gold), and EVs (magenta spheres). Dotted white line in movie shows position of the TZ. *rab-28* mutants ectopically accumulate excess EVs in distal regions of the cephalic lumen.

## REFERENCES

Akella, J.S. et al., 2019. Cell type-specific structural plasticity of the ciliary transition zone in C. elegans. Biology of the cell / under the auspices of the European Cell Biology Organization, 111(4), pp.95–107.

Ansley, S.J. et al., 2003. Basal body dysfunction is a likely cause of pleiotropic Bardet-Biedl syndrome. Nature, 425(6958), pp.628–633.

Bacaj, T. et al., 2008. Glia are essential for sensory organ function in C. elegans. Science, 322(5902), pp.744–747.

Badgandi, H.B. et al., 2017. Tubby family proteins are adapters for ciliary trafficking of integral membrane proteins. The Journal of cell biology, 216(3), pp.743–760.

Bae, Y.-K. et al., 2006. General and cell-type specific mechanisms target TRPP2/PKD-2 to cilia. Development, 133(19), pp.3859–3870.

Bargmann, C.I., 2006. Chemosensation in C. elegans. WormBook: the online review of C. elegans biology, pp.1–29.

Barr, M.M. et al., Extracellular matrix regulates morphogenesis and function of ciliated sensory organs inCaenorhabditis elegans. Available at: http://dx.doi.org/10.1101/376152.

Barr, M.M. et al., 2001. The Caenorhabditis elegans autosomal dominant polycystic kidney disease gene homologs lov-1 and pkd-2 act in the same pathway. Current biology: CB, 11(17), pp.1341– 1346.

Bhogaraju, S. et al., 2013. Molecular basis of tubulin transport within the cilium by IFT74 and IFT81. Science, 341(6149), pp.1009–1012.

Blacque, O.E., Scheidel, N. & Kuhns, S., 2018. Rab GTPases in cilium formation and function. Small GTPases, 9(1-2), pp.76–94. Available at: http://dx.doi.org/10.1080/21541248.2017.1353847.

Blanc, L. & Vidal, M., 2018. New insights into the function of Rab GTPases in the context of exosomal secretion. Small GTPases, 9(1-2), pp.95–106.

Braunreiter, K., Hamlin, S. & Lyman-Gingerich, J., 2014. Identification and characterization of a novel allele of Caenorhabditis elegans bbs-7. PloS one, 9(12), p.e113737.

Brenner, S., 1974. The genetics of Caenorhabditis elegans. Genetics, 77(1), pp.71–94.

Budnik, V., Ruiz-Cañada, C. & Wendler, F., 2016. Extracellular vesicles round off communication in the nervous system. Nature reviews. Neuroscience, 17(3), pp.160–172.

Cao, M. et al., 2015. Uni-directional ciliary membrane protein trafficking by a cytoplasmic retrograde IFT motor and ciliary ectosome shedding. eLife, 4. Available at: http://dx.doi.org/10.7554/elife.05242.

Carter, S.P. & Blacque, O.E., 2019. Membrane retrieval, recycling and release pathways that organise and sculpt the ciliary membrane. Current opinion in cell biology, 59, pp.133–139.

Cook, S.J. et al., 2019. Whole-animal connectomes of both Caenorhabditis elegans sexes. Nature, 571(7763), pp.63–71.

Diener, D.R., Lupetti, P. & Rosenbaum, J.L., 2015. Proteomic analysis of isolated ciliary transition zones reveals the presence of ESCRT proteins. Current biology: CB, 25(3), pp.379–384.

Dilan, T.L. et al., 2018. Bardet–Biedl syndrome-8 (BBS8) protein is crucial for the development of outer segments in photoreceptor neurons. Human Molecular Genetics, 27(2), pp.283–294. Available at: http://dx.doi.org/10.1093/hmg/ddx399.

Doroquez, D.B. et al., 2014. A high-resolution morphological and ultrastructural map of anterior sensory cilia and glia in Caenorhabditis elegans. eLife, 3, p.e01948.

Eguether, T. et al., 2014. IFT27 links the BBSome to IFT for maintenance of the ciliary signaling compartment. Developmental cell, 31(3), pp.279–290.

Fansa, E.K. et al., 2016. PDE6δ-mediated sorting of INPP5E into the cilium is determined by cargo-carrier affinity. Nature communications, 7, p.11366.

Fan, Y. et al., 2004. Mutations in a member of the Ras superfamily of small GTP-binding proteins causes Bardet-Biedl syndrome. Nature genetics, 36(9), pp.989–993.

Fröhlich, D. et al., 2014. Multifaceted effects of oligodendroglial exosomes on neurons: impact on neuronal firing rate, signal transduction and gene regulation. Philosophical transactions of the Royal Society of London. Series B, Biological sciences, 369(1652). Available at: http://dx.doi.org/10.1098/rstb.2013.0510.

Frühbeis, C. et al., 2013. Neurotransmitter-triggered transfer of exosomes mediates oligodendrocyte-neuron communication. PLoS biology, 11(7), p.e1001604.

Goncalves, M.B. et al., 2015. Neuronal RARβ Signaling Modulates PTEN Activity Directly in Neurons and via Exosome Transfer in Astrocytes to Prevent Glial Scar Formation and Induce Spinal Cord Regeneration. The Journal of Neuroscience, 35(47), pp.15731–15745. Available at: http://dx.doi.org/10.1523/jneurosci.1339-15.2015.

Hogan, M.C. et al., 2009. Characterization of PKD protein-positive exosome-like vesicles. Journal of the American Society of Nephrology: JASN, 20(2), pp.278–288.

Hsu, C. et al., 2010. Regulation of exosome secretion by Rab35 and its GTPase-activating proteins TBC1D10A–C. The Journal of Cell Biology, 189(2), pp.223–232. Available at: http://dx.doi.org/10.1083/jcb.200911018.

Hsu, Y. et al., 2017. BBSome function is required for both the morphogenesis and maintenance of the photoreceptor outer segment. PLoS genetics, 13(10), p.e1007057.

Ismail, S.A. et al., 2012. Structural basis for Arl3-specific release of myristoylated ciliary cargo from UNC119. The EMBO journal, 31(20), pp.4085–4094.

Jensen, V.L. et al., 2016. Whole-Organism Developmental Expression Profiling Identifies RAB-28 as a Novel Ciliary GTPase Associated with the BBSome and Intraflagellar Transport. PLoS genetics, 12(12), p.e1006469.

Jensen, V.L. & Leroux, M.R., 2017. Gates for soluble and membrane proteins, and two trafficking systems (IFT and LIFT), establish a dynamic ciliary signaling compartment. Current opinion in cell biology, 47, pp.83–91.

Jin, H. et al., 2010. The conserved Bardet-Biedl syndrome proteins assemble a coat that traffics membrane proteins to cilia. Cell, 141(7), pp.1208–1219.

Kaplan, O.I. et al., 2012. Endocytosis Genes Facilitate Protein and Membrane Transport in C. elegans Sensory Cilia. Current Biology, 22(6), pp.451–460. Available at: http://dx.doi.org/10.1016/j.cub.2012.01.060.

Knödler, A. et al., 2010. Coordination of Rab8 and Rab11 in primary ciliogenesis. Proceedings of the National Academy of Sciences of the United States of America, 107(14), pp.6346–6351.

Kösling, S.K. et al., 2018. Mechanism and dynamics of INPP5E transport into and inside the ciliary compartment. Biological chemistry, 399(3), pp.277–292.

Lechtreck, K.F., 2015. IFT-Cargo Interactions and Protein Transport in Cilia. Trends in biochemical sciences, 40(12), pp.765–778.

Lechtreck, K.-F. et al., 2009. The Chlamydomonas reinhardtii BBSome is an IFT cargo required for export of specific signaling proteins from flagella. The Journal of cell biology, 187(7), pp.1117– 1132.

Lee, G.-I. et al., 2017. A novel likely pathogenic variant in the RAB28 gene in a Korean patient with cone-rod dystrophy. Ophthalmic genetics, 38(6), pp.587–589.

Liem, K.F., Jr et al., 2012. The IFT-A complex regulates Shh signaling through cilia structure and membrane protein trafficking. The Journal of cell biology, 197(6), pp.789–800.

Liew, G.M. et al., 2014. The intraflagellar transport protein IFT27 promotes BBSome exit from cilia through the GTPase ARL6/BBS3. Developmental cell, 31(3), pp.265–278.

Liu, P. & Lechtreck, K.F., 2018. The Bardet–Biedl syndrome protein complex is an adapter expanding the cargo range of intraflagellar transport trains for ciliary export. Proceedings of the National Academy of Sciences, 115(5), pp.E934–E943. Available at: http://dx.doi.org/10.1073/pnas.1713226115.

Li, Y. & Hu, J., 2011. Small GTPases and cilia. Protein & Cell, 2(1), pp.13–25. Available at: http://dx.doi.org/10.1007/s13238-011-1004-7.

Long, H. et al., 2016. Comparative Analysis of Ciliary Membranes and Ectosomes. Current biology: CB, 26(24), pp.3327–3335.

Lopez-Verrilli, M.A., Picou, F. & Court, F.A., 2013. Schwann cell-derived exosomes enhance axonal regeneration in the peripheral nervous system. Glia, 61(11), pp.1795–1806.

Lumb, J.H. et al., 2011. Rab28 function in trypanosomes: interactions with retromer and ESCRT pathways. Journal of cell science, 124(Pt 22), pp.3771–3783.

Maas, S.L.N., Breakefield, X.O. & Weaver, A.M., 2017. Extracellular Vesicles: Unique Intercellular Delivery Vehicles. Trends in cell biology, 27(3), pp.172–188.

Maguire, J.E. et al., 2015. Myristoylated CIL-7 regulates ciliary extracellular vesicle biogenesis. Molecular biology of the cell, 26(15), pp.2823–2832.

Mangeol, P., Prevo, B. & Peterman, E.J.G., 2016. KymographClear and KymographDirect: two tools for the automated quantitative analysis of molecular and cellular dynamics using kymographs. Molecular biology of the cell, 27(12), pp.1948–1957.

Masyuk, A.I. et al., 2010. Biliary exosomes influence cholangiocyte regulatory mechanisms and proliferation through interaction with primary cilia. American journal of physiology. Gastrointestinal and liver physiology, 299(4), pp.G990–9.

Meldolesi, J., 2018. Exosomes and Ectosomes in Intercellular Communication. Current Biology, 28(8), pp.R435–R444. Available at: http://dx.doi.org/10.1016/j.cub.2018.01.059.

Molina-García, L. et al., A direct glia-to-neuron natural transdifferentiation ensures nimble sensory-motor coordination of male mating behaviour. Available at: http://dx.doi.org/10.1101/285320.

Morsci, N.S. & Barr, M.M., 2011. Kinesin-3 KLP-6 regulates intraflagellar transport in male-specific cilia of Caenorhabditis elegans. Current biology: CB, 21(14), pp.1239–1244.

Mukhopadhyay, S. et al., 2010. TULP3 bridges the IFT-A complex and membrane phosphoinositides to promote trafficking of G protein-coupled receptors into primary cilia. Genes & development, 24(19), pp.2180–2193.

Muralidharan-Chari, V. et al., 2009. ARF6-regulated shedding of tumor cell-derived plasma membrane microvesicles. Current biology: CB, 19(22), pp.1875–1885.

Nachury, M.V., 2018. The molecular machines that traffic signaling receptors into and out of cilia. Current Opinion in Cell Biology, 51, pp.124–131. Available at: http://dx.doi.org/10.1016/j.ceb.2018.03.004.

Nachury, M.V. & Mick, D.U., 2019. Establishing and regulating the composition of cilia for signal transduction. Nature reviews. Molecular cell biology. Available at: http://dx.doi.org/10.1038/s41580-019-0116-4.

Nager, A.R. et al., 2017. An Actin Network Dispatches Ciliary GPCRs into Extracellular Vesicles to Modulate Signaling. Cell, 168(1-2), pp.252–263.e14.

Nechipurenko, I.V. et al., 2017. Centriolar remodeling underlies basal body maturation during ciliogenesis in Caenorhabditis elegans. eLife, 6. Available at: http://dx.doi.org/10.7554/elife.25686.

O’Hagan, R. et al., 2017. Glutamylation Regulates Transport, Specializes Function, and Sculpts the Structure of Cilia. Current Biology, 27(22), pp.3430–3441.e6. Available at: http://dx.doi.org/10.1016/j.cub.2017.09.066.

Oikonomou, G. et al., 2011. Opposing Activities of LIT-1/NLK and DAF-6/Patched-Related Direct Sensory Compartment Morphogenesis in C. elegans. PLoS Biology, 9(8), p.e1001121. Available at: http://dx.doi.org/10.1371/journal.pbio.1001121.

Ostrowski, M. et al., 2010. Rab27a and Rab27b control different steps of the exosome secretion pathway. Nature cell biology, 12(1), pp.19–30; sup pp 1–13.

Perens, E.A. & Shaham, S., 2005. C. elegans daf-6 encodes a patched-related protein required for lumen formation. Developmental cell, 8(6), pp.893–906.

Perkins, L.A. et al., 1986. Mutant sensory cilia in the nematode Caenorhabditis elegans. Developmental biology, 117(2), pp.456–487.

Phua, S.C. et al., 2017. Dynamic Remodeling of Membrane Composition Drives Cell Cycle through Primary Cilia Excision. Cell, 168(1-2), pp.264–279.e15.

Raposo, G. & Stoorvogel, W., 2013. Extracellular vesicles: exosomes, microvesicles, and friends. The Journal of cell biology, 200(4), pp.373–383.

Reiter, J.F. & Leroux, M.R., 2017. Genes and molecular pathways underpinning ciliopathies. Nature reviews. Molecular cell biology, 18(9), pp.533–547.

Riveiro-Álvarez, R. et al., 2015. New mutations in the RAB28 gene in 2 Spanish families with cone-rod dystrophy. JAMA ophthalmology, 133(2), pp.133–139.

Roosing, S. et al., 2013. Mutations in RAB28, encoding a farnesylated small GTPase, are associated with autosomal-recessive cone-rod dystrophy. American journal of human genetics, 93(1), pp.110–117.

Rosenbaum, J.L. & Witman, G.B., 2002. Intraflagellar transport. Nature Reviews Molecular Cell Biology, 3(11), pp.813–825. Available at: http://dx.doi.org/10.1038/nrm952.

Salinas, R.Y. et al., 2017. Photoreceptor discs form through peripherin-dependent suppression of ciliary ectosome release. The Journal of cell biology, 216(5), pp.1489–1499.

Sanders, A.A.W.M., Kennedy, J. & Blacque, O.E., 2015. Image analysis of Caenorhabditis elegans ciliary transition zone structure, ultrastructure, molecular composition, and function. Methods in cell biology, 127, pp.323–347.

Schindelin, J. et al., 2012. Fiji: an open-source platform for biological-image analysis. Nature methods, 9(7), pp.676–682.

Serwas, D. et al., 2017. Centrioles initiate cilia assembly but are dispensable for maturation and maintenance in C. elegans. The Journal of Cell Biology, 216(6), pp.1659–1671. Available at: http://dx.doi.org/10.1083/jcb.201610070.

Silva, M. et al., 2017. Cell-Specific α-Tubulin Isotype Regulates Ciliary Microtubule Ultrastructure, Intraflagellar Transport, and Extracellular Vesicle Biology. Current biology: CB, 27(7), pp.968– 980.

Sulston, J.E., Albertson, D.G. & Thomson, J.N., 1980. The Caenorhabditis elegans male: Postembryonic development of nongonadal structures. Developmental biology, 78(2), pp.542– 576.

Wang, J. et al., 2014. C. elegans Ciliated Sensory Neurons Release Extracellular Vesicles that Function in Animal Communication. Current Biology, 24(5), pp.519–525. Available at: http://dx.doi.org/10.1016/j.cub.2014.01.002.

Wang, J. et al., 2015. Cell-Specific Transcriptional Profiling of Ciliated Sensory Neurons Reveals Regulators of Behavior and Extracellular Vesicle Biogenesis. Current biology: CB, 25(24), pp.3232–3238.

Wang, J. & Barr, M.M., 2018. Cell-cell communication via ciliary extracellular vesicles: clues from model systems. Essays in biochemistry, 62(2), pp.205–213.

Wang, T. et al., 2014. Hypoxia-inducible factors and RAB22A mediate formation of microvesicles that stimulate breast cancer invasion and metastasis. Proceedings of the National Academy of Sciences of the United States of America, 111(31), pp.E3234–42.

Ward, S. et al., 1975. Electron microscopical reconstruction of the anterior sensory anatomy of the nematodecaenorhabditis elegans. The Journal of Comparative Neurology, 160(3), pp.313–337. Available at: http://dx.doi.org/10.1002/cne.901600305.

Waters, A.M. & Beales, P.L., 2011. Ciliopathies: an expanding disease spectrum. Pediatric Nephrology, 26(7), pp.1039–1056. Available at: http://dx.doi.org/10.1007/s00467-010-1731-7.

Wehman, A.M. et al., 2011. The P4-ATPase TAT-5 inhibits the budding of extracellular vesicles in C. elegans embryos. Current biology: CB, 21(23), pp.1951–1959.

Weimer, R.M., 2006. Preservation of *C. elegans* Tissue Via High-Pressure Freezing and Freeze-Substitution for Ultrastructural Analysis and Immunocytochemistry. In S. Kevin, ed. C. elegans. New Jersey: Humana Press, pp. 203–222.

Wood, C.R. et al., 2013. The cilium secretes bioactive ectosomes. Current biology: CB, 23(10), pp.906–911.

Xu, Q. et al., 2015. BBS4 and BBS5 show functional redundancy in the BBSome to regulate the degradative sorting of ciliary sensory receptors. Scientific reports, 5, p.11855.

Ye, F., Nager, A.R. & Nachury, M.V., 2018. BBSome trains remove activated GPCRs from cilia by enabling passage through the transition zone. The Journal of cell biology, 217(5), pp.1847–1868.

Ying, G. et al., 2018. The small GTPase RAB28 is required for phagocytosis of cone outer segments by the murine retinal pigmented epithelium. The Journal of biological chemistry, 293(45), pp.17546–17558.

Zappulli, V. et al., 2016. Extracellular vesicles and intercellular communication within the nervous system. Journal of Clinical Investigation, 126(4), pp.1198–1207. Available at: http://dx.doi.org/10.1172/jci81134.

